# High resolution spatial transcriptomic and proteomic profiling of early primate gastrulation *in utero*

**DOI:** 10.64898/2026.02.10.705168

**Authors:** Nikola Sekulovski, Maliha Kabir, Anusha Rengarajan, Amber E. Carleton, Jenna K. Schmidt, Chien-Wei Lin, Kenichiro Taniguchi

## Abstract

Early gastrulation represents a key stage in which several embryonic and extra-embryonic lineages are formed in the primitive streak to support embryonic development. However, very little is known about lineage specification events in the early primate gastrula. To gain molecular insights into mechanisms that organize this stage of primate development, we performed high-resolution spatial transcript and protein expression profiling of five sagittal sections from a cynomolgus macaque embryo at Carnegie Stage 6b, an early gastrulation stage. We established a molecular map of six major cell populations: the epiblast, primitive streak, amnion, endoderm, mesoderm, and primordial germ cells. We also uncovered a variety of lineage subtypes, as well as important signaling and transcriptional networks. In particular, we show that canonical WNT signaling is a driver of amnion fate progression. Together, this study provides a unique multiomic resource of an early primate gastrula with complete spatial information for future investigations.

## Introduction

Immediately prior to gastrulation, the primate embryo is composed of four major cell types: epiblast, amnion, extra-embryonic endoderm (ExEE) and trophectoderm [1–3]. The onset of gastrulation is marked by formation of the primitive streak (PrSt) at the posterior tip of the epiblast; the PrSt will give rise to definitive endoderm (DE), mesoderm, primordial germ cells (PGC) and amnion [3–5]. Studies in mice first showed that PrSt positioning is controlled by the visceral endoderm (VE, also referred to as the hypoblast), a component of ExEE [3, 5]. Mechanistically, the VE tissue that underlies the nascent anterior domain of the epiblast (called the anterior visceral endoderm, AVE) expresses secreted inhibitors of several signaling pathways (TGFβ/Nodal, BMP, and WNT) that are critical for inducing PrSt formation, thereby posteriorly restricting PrSt formation [6, 7]. Several recent studies now show that the AVE from early primate embryos also expresses such secreted signaling inhibitors [8–10]. Importantly, the amnion was recently shown to serve as a signaling hub to support mesoderm induction in primates [3, 11, 12]. Therefore, crosstalk between these early embryonic and extra-embryonic cell types is critical for supporting early embryogenesis.

In the past decade, two landmark single cell transcriptomic studies have advanced our understanding of early primate development *in situ* [13, 14]. Additionally, advances in spatial transcriptomic technology [15–18] are providing unprecedented insights into early primate embryology. A study by Bergmann *et al.* used a laser capture microdissection approach on *in utero* marmoset embryos staged at Carnegie stage 5 (CS5, late implantation/pre-gastrulation), CS6 (early gastrulation) or CS7 (mid-late gastrulation), achieving spatial profiling of representative spots with 20-30μm diameter (**Fig. S1A**) with gaps of at least 50μm between spots (**Fig. S1B**) [9]. Additionally, two recent studies by Cui *et al.* [19] and Xiao *et al.* [20] explored the transcriptome of human embryos staged at CS7 and CS8 (neural plate stage), respectively. Using a Stereo-seq platform [21], these authors systematically profiled transcriptomes from an array of circular 220nm diameter DNA nanoballs with 500nm-715nm center-to-center spacing. To obtain sufficient sequencing depth, an area from 50 x 50 nanoballs was then combined into bins that each cover, on average, 882μm^2^ in area (∼29.7μm x 29.7μm) (**Fig. S1A-C**) [19, 20]. Though valuable, these studies present relatively low-level resolution analyses (**Fig. S1D,E**). Here, we used a multiomic approach to molecularly profile an early primate gastrula *in utero*, resulting in single cell-level spatial resolution.

Using Visium HD (10X Genomics), a spatial transcriptomic analysis platform, and a multiplex-immunofluorescent (IF) method (called 4i) [22] that we modified for formalin fixed paraffin embedded (FFPE) tissue samples (thereafter referred to as the FFPE-4i) [23], we mapped transcript and protein expression output on five serial sagittal slices from an intact cynomolgus macaque (*Macaca fascicularis*) embryo staged at CS6b. The spatial transcriptomics approach resulted in an improved resolution (bins per 100×100 μm^2^) by 78-fold (compared to Bergmann *et al.*) and 14-fold (compared to Cui *et al.* and Xiao *et al.*). Moreover, our spatial proteomic approach achieved spatial profiling at the single cell level. Using this high-resolution multiomic resource, we molecularly defined subtypes of cell lineages in the early primate gastrula, and uncovered previously unrecognized aspects of lineage specification and morphogenesis during early primate development.

## Results

### Transcriptomic and proteomic pipelines to spatially profile a Carnegie stage 6b early cynomolgus macaque gastrula

The probe-based Visium HD platform enables robust high-resolution spatial transcriptomic profiling with 4μm^2^ resolution (2μm x 2μm square tiles, **Fig. S1A**) without spacing (**Fig. S1B**); sixteen of the 2μm tiles are combined, which covers 64μm^2^ in area (thereafter referred to as 8μm bin, **Fig. S1C**), to obtain sufficient sequencing depth [18]. Therefore, the Visium HD offers a spatially complete (gap-less) platform, which represents a significant improvement in resolution compared to earlier studies [9, 19, 20] (**Fig S1A-E**) with approximately single cell resolution per bin based on our deconvolution analysis using STdeconvolve [24] (**Fig. S1F**). Recently, we successfully conducted a multiplexed (14-plex) protein expression analysis in early cynomolgus macaque embryo section samples using FFPE-4i, presenting a multiplexed IF pipeline to investigate early primate embryo samples at single cell levels [23]. In this study, using the Visium HD platform (with probes designed for human transcripts) and the FFPE-4i pipeline, we explore the spatial transcriptome and proteome of FFPE sections from a separate early cynomolgus macaque embryo.

This embryo, with an approximate gestation date of 16-17 based on our ultrasound analysis of the gestational sac (**Fig. S1G**) was identified in serial transverse sections (5μm) of FFPE uterine tissue from a pregnant cynomolgus macaque female. The amnion, the embryonic disc with a convex morphology, and a small amniotic lumen are readily seen, while a clear prechordal plate– or node-like structure is not seen (**Fig. 1A**). Importantly, the edge to edge diameter of the embryonic disc is ∼250μm, and the embryo dimension is ∼350μm x ∼280μm (**Fig. 1A**). Together, the approximate gestation date, overall embryo morphology and measurements estimate this embryo at CS6.

**Figure 1.**
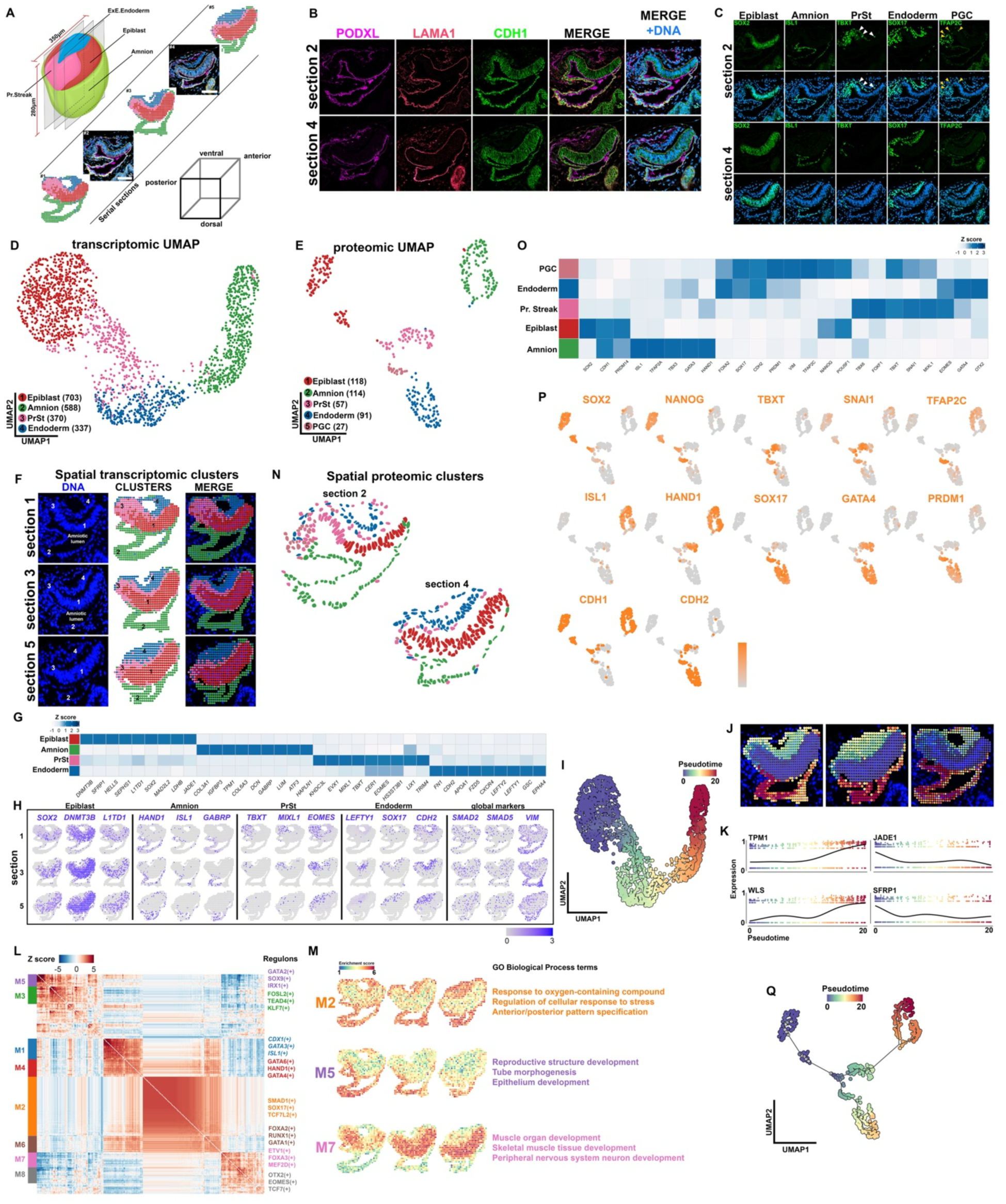
Spatial transcriptomic and proteomic profiling of a natural CS6b cynomolgus macaque embryo. **A**) A schematic representation for spatial profiling of the CS6b embryo using transcriptomic (sections #1, 3, and 5) and proteomic (sections #2 and 4) approaches. The gray cube is used to orient the embryonic axes. Scale bar, 100µm. **B,C)** Confocal images of immuno-stained sections of the CS6b embryo using indicated polarized epithelium (B) and lineage (C) markers. **D)** A UMAP plot displaying the coordinates of each spatial transcriptome bin with four identified populations: 1 – Epiblast (red), 2 – Amnion (Green), 3 – Primitive Streak (pink), 4 – Endoderm (blue). Numbers represent bins per cluster. **E)** A UMAP plot displaying the coordinates of each cell from the spatial proteomic analysis with five identified populations: 1 – Epiblast (red), 2 – Amnion (green), 3 – Primitive Streak (pink), 4 – Endoderm (blue), 5 – PGC (coral). Numbers represent cells per cluster. **F)** Spatial distribution of the transcriptomic bins labeled with identified populations in (D). **G)** Heatmap based on the normalized expression values from the transcriptional dataset of top 9 differentially enriched genes in each cell lineage. **H)** Expression of selected lineage specific and globally expressed genes superimposed onto the spatial transcriptome plot. **I)** Pseudotime analysis of the transcriptomic dataset, showing predicted lineage progression trajectories, based on Monocle3. **J)** Spatial distribution of lineage progression trajectories based on transcriptomic pseudotime analysis in (I). **K)** Scatterplots showing the expression dynamics of indicated genes along the pseudotime trajectory in (J). **L,M)** Two-dimensional heatmap of gene regulatory networks based on the SCENIC analysis, which are grouped based on spatial correlation using the Hotspot package, identifying eight modules (M1-M8, see (M) for spatially projected enrichment scores for M2, M5 and M7 with selected GO Biological Processes terms, also see **Fig. S4D** for all modules). **N)** Spatial projection of cell lineages based on (E). Nucleus from each quantitated cell in the proteomic dataset are labeled with cell lineage. **O)** Heatmap based on the normalized signal intensity from the proteomics dataset of selected lineage-specific transcription factors. **P)** Expression of indicated lineage-specific factors superimposed onto the spatial proteomics UMAP plot in (E). **Q)** Pseudotime analysis of the proteomics dataset, showing predicted lineage progression trajectories that are similar to the transcriptomic dataset.

To gain molecular insight into cell identities, spatial organization and gene expression during early primate gastrulation, a multiomic analysis was performed using five sagitally-oriented serial sections (**Fig. 1A**). Sections 1, 3 and 5 were used to uncover the spatial transcriptome using the Visium HD platform, and sections 2 and 4 were processed to explore the spatial proteome using the FFPE-4i pipeline (**Fig. 1A**). This approach, which positions transcriptomics and proteomics in alternating sections, allows us to robustly measure transcriptomic and proteomic relationships at spatial levels.

Using Visium HD, an area of 42.25mm^2^ (6.5mm x 6.5mm), containing the embryo, placenta and uterus was examined in total (**Fig. S1H-J**), obtaining 11,235,904 (3,352 x 3,352) 2μm tiles, which were combined into 702,244 (838 x 838) 8μm bins for downstream analyses (profiling 2,500 tiles or 156 bins per 10^4^μm^2^ area, **Fig. S1D,E**). Given that human and cynomolgus macaque genomes share more than 93% similarity [25–27], the Visium HD human probes can identify 15,222 cynomolgus macaque protein coding genes (see **Table S1** for the manufacturer-provided list of detectable genes). Followed by alignment, demultiplexing and spatial assignment of the reads using Space Ranger (v3.0), we captured an average of 181.6 UMI (unique molecular identifier) across all 8μm bins in the entire scanned area, including the uterine cavity. In this study, we examine spatial transcriptomic profiles of embryo-derived, but not placental, components (**Fig. 1A**, see dotted boxes in **Fig. S1H-J**), which show an average of 695 UMI and 607 features per bin across 1,998 total 8μm bins (744,690 and 564 bins in sections 1, 3 and 5, respectively), totaling 108,420 UMI per 10^4^μm^2^ area (**Fig. S1K-M**). These output parameters are similar to those obtained in previous spatial transcriptomic studies based on Visium HD using human samples [16, 18].

Using the FFPE-4i pipeline (**Fig. S1N**, 40 total cycles) for slides 2 and 4, we achieved multiplexed labeling of 61 unique proteins. For each IF cycle, tiled confocal images covering 1.82mm^2^ (1.35mm x 1.35mm) in area were obtained, which were then quantitated for expression intensity per cell (nucleus for 36 transcription factors and two proliferation markers, binary for 3 cell membrane, 1 cytoplasmic, and 3 secreted proteins, **Table S2**) from embryo-derived, but not placental, components (section 2 – 216 cells; section 4 – 191 cells, **Fig. S1O,** selected markers shown in **Fig. 1B,C**, **S1P**; all markers shown in **Fig. S2A,B**). Additional 9 cell membrane (e.g., PODXL, LAMA1, ZO-1) and 4 intracellular (e.g., RAB5, RAB7, RAB11) markers were used to show previously unrecognized cell biological characteristics of early embryonic lineages (**Fig. S2A,B**). Importantly, at least one marker per known cell lineage was included in more than one IF cycle as controls to test for protein degradation and/or reduced antigen detectability; these markers show similar expression intensity in each IF cycle (see **Fig. S2C** for the replicate reproducibility analysis based on hierarchical clustering, adjusted Rand index (ARI = 1)).

Embryo structure was next examined using the FFPE-4i dataset. Localization of PODXL and LAMA1 shows the formation of apical and basolateral membrane territories, respectively (**Fig. 1B**). Moreover, CDH1 (E-Cadherin), CDH2 (N-Cadherin) and TJP1 (ZO-1) display highly organized membrane localization in several tissues including the epiblast, amnion and ExEE, which are marked by SOX2, ISL1 and SOX17, respectively (**Fig. 1B-C, S1P**). The epiblast tissue in the section 2 (medial) is overall thinner than that tissue in the section 4 (slightly more lateral, **Fig. S2D**), which is likely due to sectioning the convex epiblast structure. Restricted TBXT staining is seen on one side of the epiblast in both sections, revealing the formation of the primitive streak (**Fig. 1C**). Also, several disseminated TBXT^+^ cells are present between the ExEE and the epiblast (**Fig. 1C**, white arrowheads), which were not seen at CS6a [23]. These results show well-preserved embryonic structures, and provide positional information for the anterior-posterior (A-P) axis (**Fig. 1A**), as well as additional molecular details to stage this embryo at CS6b, the early-mid gastrulation stage.

Together, these complementary molecular approaches present a high-resolution spatial multiomic map during early primate gastrulation (**Fig. 1A**, coordinates of each bin (transcriptome) or cell (proteome) in a dimensionally-reduced space as shown in **Fig. 1D,E** with cell type annotation (further described below)).

### Global analysis of the spatial transcriptomic dataset

The transcriptomic dataset was examined using Seurat [28] for data filtering, normalization, variable gene selection and subsequent unsupervised clustering of bins (see Materials and Methods, **Table S3**). The transcriptomic datasets from the three sections were integrated and the data yielded four distinct populations among the 1,998 total 8μm bins (**Fig. 1D, S3A**), which, after marker and lineage analyses described below, are labeled Epiblast (red), Amnion (green), Primitive Streak (PrSt, pink) and Endoderm (blue).

Consistent with the positions of the four clusters in the embryo, differential gene expression analysis shows enrichment of known markers of the Epiblast (expressing *SOX2*, as well as *DNMT3B*, and *L1TD1* [14, 19, 20], Amnion (*COL3A1*, *IGFBP3*, *GABRP)* [12, 14, 20, 29], PrSt (*MIXL1, TBX1*, *EOMES*) [14, 30, 31], and Endoderm (*CDH2*, *LEFTY1*, *GSC)* [9, 11], (**Fig. 1F,G, Table S4**, additional genes are shown in **S3B,C**), demonstrating the high fidelity of our dataset at molecular levels. Notably, in addition to genes that show expression pattern that is specific to one cell type (e.g., *SOX2*, *GABRP*, *LEFTY1*), several genes also show known expression in more than one lineage (e.g., *ISL1* and *HAND1* (amnion and mesoderm), *GSC* (PrSt and endoderm)), **Fig. 1H**), revealing that our dataset is highly resolved. Importantly, trajectory analysis using the Monocle 3 package [32] shows predicted lineage progression during early primate development (**Fig. 1I, S4A**, see **Fig. 1J** for spatial projection, see **Fig. 1K, S4B** for selected differentially expressed genes (DEGs, from **Fig. 1G**, **Table S4**) with dynamic expression along the lineage trajectory).

We next explored transcriptional regulatory networks based on the SCENIC (single-cell regulatory inference and clustering) [33] package. The SCENIC analysis shows enriched activity of several transcriptional regulatory networks with known links to specific lineages (e.g., Epiblast: MAFK, MYC; Amnion: HAND1, GATA3; PrSt: EOMES, CDX1; endoderm: GATA4, FOXA1), spatially defining tissue-specific transcriptional networks (**Fig. S4C**, **Table S5**). Indeed, regulon enrichment analysis based on Hotspot [34] identifies 8 major tissue specific modules (**Table S6**), spatially defining several tissue-specific transcriptional network activities with known biological functions based on Gene Ontology (**Fig. 1L,M, S4D**, **Table S7**).

These results show that this dataset provides a high-resolution transcriptomic map of early primate gastrulation, which can be further examined using a variety of computational pipelines for investigations into active transcriptional activities across early primate embryonic lineages.

### Global analysis of the spatial proteomic dataset

Data normalization, variable feature selection and subsequent unsupervised clustering of the spatial proteomic dataset was performed by the Seurat pipeline (see Materials and Methods), which yielded five distinct populations among 407 total cells across two sections, and all five cell lineages map to corresponding territories in the embryo (**Fig. 1N)**. In addition to the four major populations that are seen in the spatial transcriptomic dataset (Epiblast (red, SOX2 and NANOG), Amnion (green, HAND1 and ISL1), PrSt (pink, TBXT and SNAI1) and endoderm (blue, SOX17 and GATA4), we also identified PGC (coral, TFAP2C, NANOG and SOX17, **Fig. 1O,P**). Moreover, trajectory analysis based on pseudotime shows predicted cluster relationships that are consistent with known lineage progressions during gastrulation (**Fig. 1Q, S4E,F**). These results show that the proteomic dataset robustly captures the molecular dynamics of the primate gastrula, and complements the transcriptomic dataset.

Also, this spatial proteomic dataset enables us to explore molecular aspects of the primate peri-gastrula at subcellular and tissue levels. Similar to our previous findings using the CS6a embryo [23], while CDH1 expression is concentrated in the epiblast and amnion, CDH2 expression is enriched at the PrSt and endoderm territories (**Fig. 1B,P, S2A,B**). In the same study [23], we also showed that PGC, marked by TFAP2C, NANOG, SOX17 and PRDM1, are seen within the PrSt territory of the CS6a embryo. Interestingly, in the CS6b embryo from this study, several PGC are adjacent to the ExEE territory, while remaining PGC are in the PrSt territory (**Fig. 1C** (yellow arrowheads)**,N,O,P)**revealing that PGC migration into the ExEE territory is initiated by CS6b. PGC and endoderm are also closely clustered in a dimensionally reduced space (**Fig. 1E**). These results reveal spatial and proteomic similarities between PGC and endoderm, and the proteomic similarity provides a reasonable explanation for how PGC may be incorporated into the endoderm territory during early development [8, 35, 36].

Together, the spatial proteomic dataset is a true single cell level resource that accompanies the single cell-like spatial transcriptomic dataset, and enables us to uncover previously unrecognized protein expression dynamics at subcellular levels during early primate gastrulation.

### Global multiomic analysis

To gain insights at multiomic levels, we next integrated the two datasets into one multiomic dataset using canonical correlation analysis [37]. Integration analysis of the two datasets shows that corresponding cell populations from each dataset indeed overlap (**Fig. 2A-C, S5A-D,** see **Fig. S6** for spatial comparisons of transcript and protein expression), revealing that transcript/protein expression dynamics are globally similar between the two datasets (**Fig. 2D**). Moreover, the pseudotime trajectory analysis also shows known lineage progressions (**Fig. 2E, S5B,C**). Our previous study, using a stem cell-derived model, showed that cells with an early primitive streak-like transcriptome emerge before the amnion is formed [23]; these lineage characteristics are also seen in this multiomics trajectory, with PrSt cells bifurcating into endoderm and amnion lineages (**Fig. 2B,C,E, S5B**), supporting this notion in an intact early primate gastrula sample.

**Figure 2.**
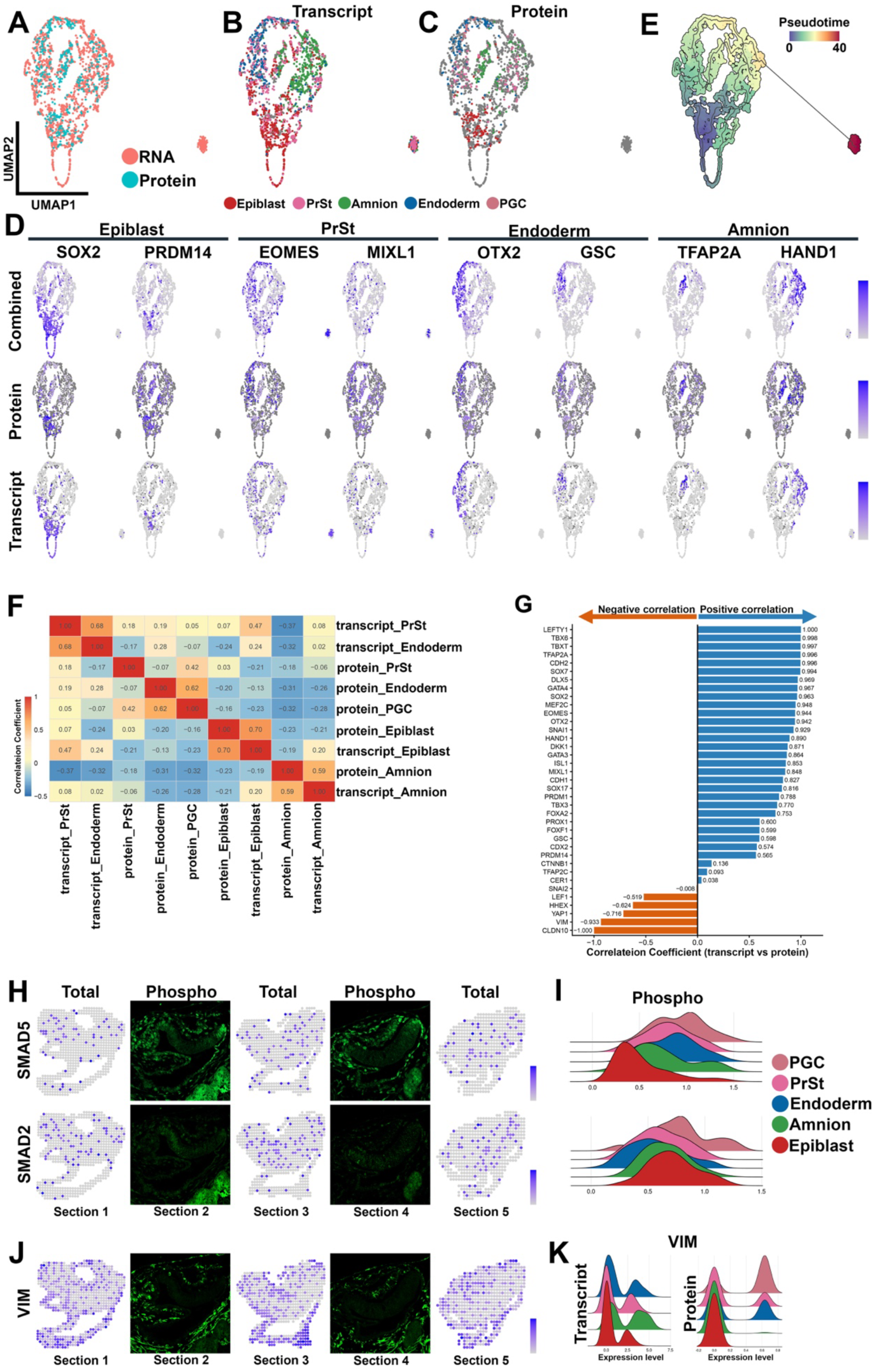
Multiomic-level analysis of the integrated transcriptomic and proteomic dataset. **A-C**) Multiomic UMAP plots showing distributions of the bins (transcriptome, salmon colored dots in A) and cells (proteome, light blue dots in A), as well as distributions of the annotated cell types from the transcriptomic (B) and proteomic (C) datasets. **D)** Transcript, protein, or combined (transcript and protein) expression of selected markers superimposed onto the multiomic UMAP plot in (A). **E)** Pseudotime analysis of the multiomic dataset, showing predicted lineage progression trajectories. **F)** A heatmap showing correlation coefficient values for the correlation among each of the clusters in the transcriptomic and proteomic datasets, based on hierarchical clustering analysis with complete linkage agglomeration method. **G)** A bar graph showing Pearson correlation coefficient values (X-axis) for the correlation between transcript and protein expression levels of indicated genes (Y-axis). **H-K)** Spatial-level expression of SMAD5 (H), SMAD2 (H) and VIM (J) at transcript (left, bins) and protein (right, confocal images) levels. Note that two antibodies that recognize phosphorylated forms of SMAD2 or SMAD1/5 are used in (H, phospho). Ridge plots for quantitation of pSMAD1/5 (I), pSMAD2 (I) and VIM (K) for each lineage.

Next, to further explore transcript and protein dynamics, we examined the correlation between transcript and protein expression levels across all cell lineages at cluster (**Fig. 2F**) as well as at transcript/protein (**Fig. 2G,S5D**) levels. Consistent with previous studies [38, 39], SMAD1/5 and SMAD2/3 transcripts, which encode nuclear transducers of BMP and TGFβ/Nodal signaling, respectively, are expressed ubiquitously (**Fig. 1H**, **2H, S6**). However, nuclear localization of phosphorylated forms of SMAD1/5 and SMAD2/3, readouts of BMP and TGFβ/Nodal signaling, respectively, are seen in specific cell populations (e.g., pSMAD1/5 – Amnion; pSMAD2/3 – PrSt, **Fig. 2H,I**), identifying lineage specific signaling characteristics. Strikingly, the transcript and protein levels of VIM (Vimentin), a type III intermediate filament [40, 41], are highly negatively correlated (**Fig. 2G**). In detail, while *VIM* transcript is abundantly expressed in all cell types, very little VIM protein is seen in the epiblast and amnion (**Fig. 2J,K, S2**). Moreover, VIM protein is expressed in the Endoderm (ExEE based on the proteomic data, **Fig. 2J,K, S2**), a surprising finding given that VIM is primarily expressed in mesenchymal cell types [11, 42, 43]. Thus, this multiomic comparison suggests the presence of novel mechanisms in some embryonic lineages that actively reduce VIM protein (**Fig. 2G, S5D**).

### Molecular characteristics of the epiblast, primitive streak, mesoderm and nascent definitive endoderm cells

To explore lineage progression during early gastrulation, we performed detailed analyses of the Epiblast and PrSt populations in the transcriptomic as well as in the proteomic datasets (**Fig. 3A-D**). Clustering of the two populations leads to the identification of the epiblast (Epiblast-1, sage; Epiblast-2, orange; Epiblast-3, light purple), PrSt (maroon) and mesoderm (light pink) cell populations, based on the marker analyses described below (**Fig. 3A-L, S7A,B**, **Table S8**).

**Figure 3.**
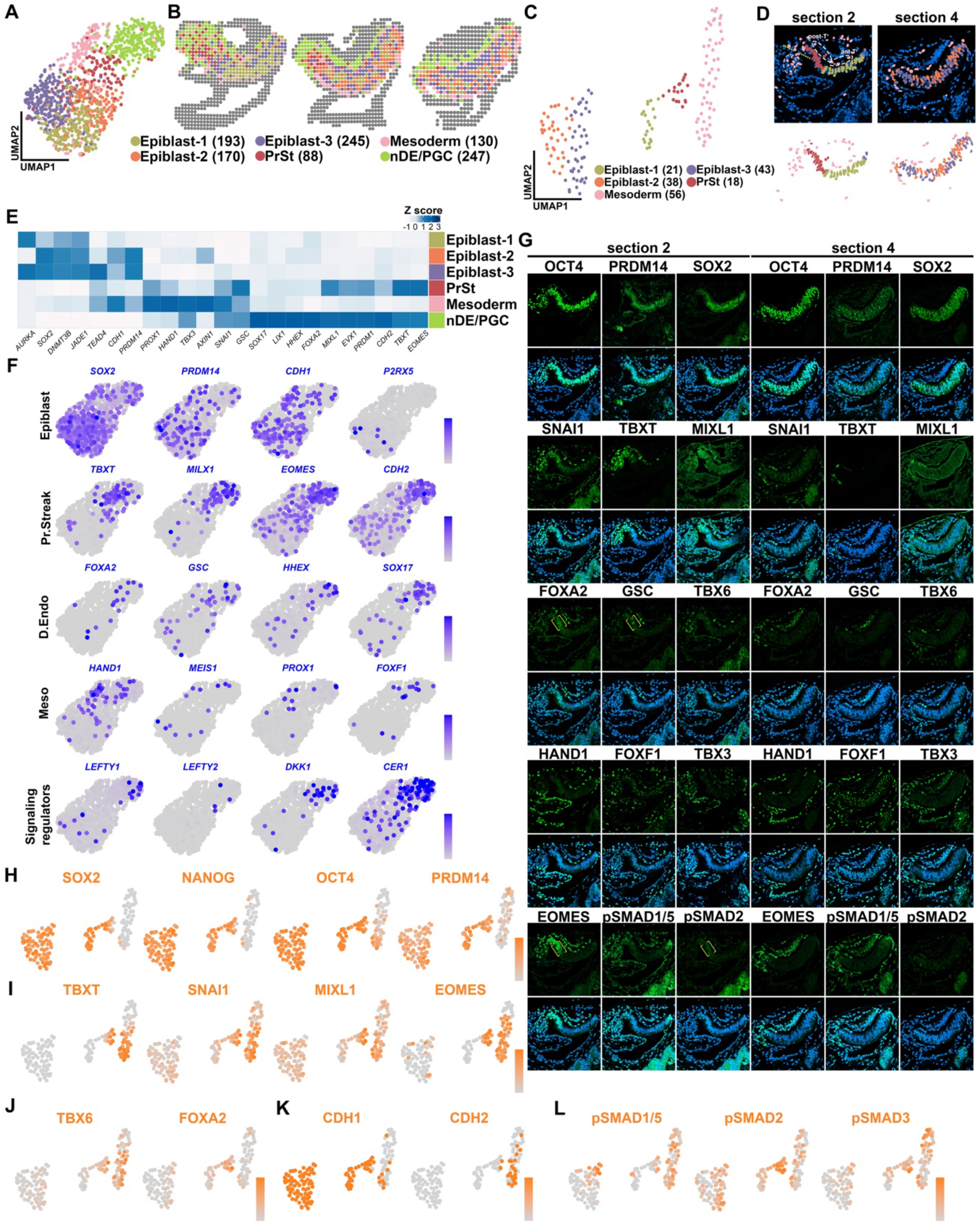
Lineage formation in the primitive streak cells. **A-D**) Sub-clustering analyses of the epiblast and PrSt lineages. Transcriptomic and proteomic UMAP plots (A and C, respectively), displaying coordinates of each bin or cell with identified populations: Epiblast-1 (sage), Epiblast-2 (orange), Epiblast-3 (light purple), PrSt (maroon), Mesoderm (light pink), nDE/PGC (lime green), which are mapped spatially in (B, transcriptomic) and (D, proteomic). Numbers represent bins/cells per cluster. Dotted yellow line in section 2 represents the posterior embryonic-extra-embryonic boundary. Pink cells in the light gray dotted circle indicates an extra-embryonic cell population with hematopoietic progenitors, allantoic progenitors and PGC. White dotted circles indicate the posterior TBXT^+^ (post-T^+^) and anterior TBXT^+^ (ant-T^+^) cell populations. **E)** A heatmap showing the normalized transcript expression values of selected lineage enriched genes. **F)** Expression of selected lineage specific genes superimposed onto the spatial transcriptomic UMAP plot (A). **G)** Confocal IF images of selected lineage markers from the proteomic dataset. Yellow brackets indicate anterior PrSt cells that are positive for FOXA2, GSC, EOMES and pSMAD2. **H-L)** Expression of known lineage-specific factors: pluripotency (H), PrSt (I), mesoderm and endoderm (J), cell-cell adhesion proteins (K), nuclear pSMAD activity (L), superimposed onto the proteomic UMAP plot in (C).

In the transcriptomic dataset, we also identify an additional cluster that contains nascent definitive endoderm (nDE) and PGC (nDE/PGC, lime green) populations, two cell lineages with a similar gene expression pattern [23, 44–46]. In the proteomic dataset, nDE cells are found in the endoderm cluster that also contains the ExEE cells (**Fig. 1O**, further described below).

Differential gene expression analyses of the transcriptomic dataset show genes that are enriched in each of the three epiblast sub-clusters (**Fig. 3E**, see **Fig. S7A** for top five DEGs). These genes are expressed in all three Epiblast subclusters, but at different levels. Interestingly, Epiblast-1 cells are seen in the medial epiblast cells that are shorter in height (sections 1 and 2) while Epiblast-2 and Epiblast-3 cells are primarily seen in the further lateral sections (sections 3-5) and display taller epiblast morphology (**Fig. 3B,D**, height quantification in **Fig. S2D**).

Furthermore, Epiblast-2 and Epiblast-3 populations show an alternating pattern along the A-P axis (purple cells at the most anterior tip, then orange, then purple) in the proteomic dataset (section 4, **Fig. 3D**); a similar spatial patterning is also observed between the Epiblast-2 and –3 populations at transcriptomic levels (p-value < 0.002, based on the spatial pairwise neighborhood analysis (see Materials and Methods)). These results show that the epiblast displays molecularly unique compartments during early gastrulation.

The PrSt (maroon) expresses known PrSt markers including TBXT and MIXL1 (**Fig. 3E-G,I**). Indeed, the PrSt population shows disorganized basal LAMA1, a key feature of epithelial-to-mesenchymal transition [47, 48], at the posterior of the epiblast that bridges the epiblast and amnion (indicated by a bracket, **Fig. S7C**). This LAMA1 disorganization also marks the embryonic-extraembryonic border (see yellow dotted line in **Fig. 3D**). Consistent with the known signaling characteristics, pSMAD2 is enriched in the nucleus of the PrSt cells (**Fig. 3G**, yellow brackets), while nuclearly localized pSMAD1/5 is spatially restricted to the PrSt cells at the epiblast-amnion boundary (**Fig. 3G,L**, see **Fig. S7D,E** for the expression of selected TGFβ/Nodal/Activin and BMP target genes).

The mesoderm (light pink) expresses known mesodermal genes such as *TBX3*, *HAND1* and *PROX1* [49–51] (**Fig. 3E,F, S7B,E**). Interestingly, TBXT protein expression is seen in all light pink mesoderm cells including those that are seen extraembryonically (indicated by a gray dotted circle in **Fig. 3D**), revealing that, similar to mice [52, 53], the T^+^ PrSt cells in primates also extend posteriorly beyond the embryo proper and into the extraembryonic territory. In mice, progenitors of hematopoietic and allantoic (referred to as body stalk in primates) lineages, as well as PGC are seen in the developing extraembryonic PrSt cells [44–46, 54]. This is also seen in the CS6b embryo examined here, based on the marker expression of hematopoietic progenitor (*MESP1* [55, 56], **Fig. S7E**), PGC (**Fig. 1C,N,O,P**), allantois (e.g., *ITGA5, HAS2* and *VCAN)* [57–59], **Fig. S7E**) and T^+^ allantoic core domain (CDH1 and TJP1, [53, 60], **Fig. S1P, S7F**) populations. Lastly, while only weak nuclear pSMAD2 expression is seen, abundant nuclear pSMAD1/5 signal and the expression of BMP target genes are observed in the mesoderm cells (**Fig. 3G,L**, **Fig. S7D**), further supporting the notion that mesoderm formation requires high BMP, but not TGFβ/Nodal/Activin, signaling activity [5, 61, 62]. These results provide spatially-resolved evidence for the presence of an extraembryonic PrSt population, as well as evidence for the BMP-driven formation of the mesoderm cells by CS6b.

In a cynomolgus macaque gastrula staged at CS6a, clear membranous CDH1 expression is seen in the PrSt cells that bridge the epiblast and amnion, while CDH2 is abundantly expressed in the mesoderm and PGC as well as in the ExEE cells [23]. Interestingly in the CS6b embryo, CDH2 protein expression is additionally seen in the layer of TBXT^+^ cells between the epiblast and ExEE (white arrowheads in **Fig. 1C**, also see **Fig. S7F**). In fact, the anteriorly localized TBXT^+^ cells with dispersed morphology (ant-T^+^, **Fig. 3D**) only express CDH2, while CDH1 expression is also seen in the posterior half of the TBXT^+^ cells (post-T^+^, **Fig. S7F**) that are more clustered (**Fig. 3G**). This result reveals the presence of two molecularly distinct TBXT^+^ cell types along the A-P axis.

Clustering analysis shows that the ant-T^+^ cells are grouped into the mesoderm population, and show enriched expression of mesoderm markers such as SNAI1, FOXF1, HAND1, TBX3 and TBX6 (**Fig. 3G, S7G**). Compared to the medial sections (sections 1 and 2), the further lateral sections (sections 3-5) show limited mesodermal cells in the anterior (**Fig. 3B,D**). Importantly, these cells do not express GSC, a marker of the established axial mesendoderm [63–65] (**Fig. 3G, S7G**). Based on these results, we suspect that the ant-T^+^ cells may represent a nascent anterior medial mesoderm (nAMM) cell population (**Fig. 3D)**.

The post-T^+^ cells are clustered within the endoderm population in the proteomic dataset (**Fig. 1N**); transcriptomic bins across this domain also show PGC/endodermal molecular signatures (**Fig. 3B**). Indeed, these CDH2^+^ post-T^+^ cells express FOXA2, CDX2 and SOX17, key markers of nDE at transcriptomic and proteomic levels (**Fig. 3E-G, S2A,B, S7G**). Moreover, these cells show SNAI1 expression levels that are lower than those seen in the ant-T^+^ cells (**Fig. S7G**), a molecular feature that is also seen in murine nDE [42]. Moreover, detailed analysis of FOXA2, GSC and EOMES expression shows that these proteins are detected in the anterior PrSt cells that are adjacent to the nDE cells (**Fig. 3G**, yellow brackets). Together, we present spatial and molecular evidence for the presence of the nAMM and nDE cells during early gastrulation, as well as for the notion that, similar to mice [66, 67], nDE cells are specified in the anterior PrSt.

### Molecular characteristics of the ExEE and amnion

To explore molecularly distinct ExEE compartments, we performed clustering analyses of the Endoderm population in the transcriptomic and proteomic datasets, which identified three cell types that spatially and molecularly match VE (light blue), AVE (blue) and ParE (parietal endoderm, cyan) (**Fig. 4A-D**, also see **Fig. 4E,F, S8A** (top 10 DEGs), **Table S9**).

**Figure 4.**
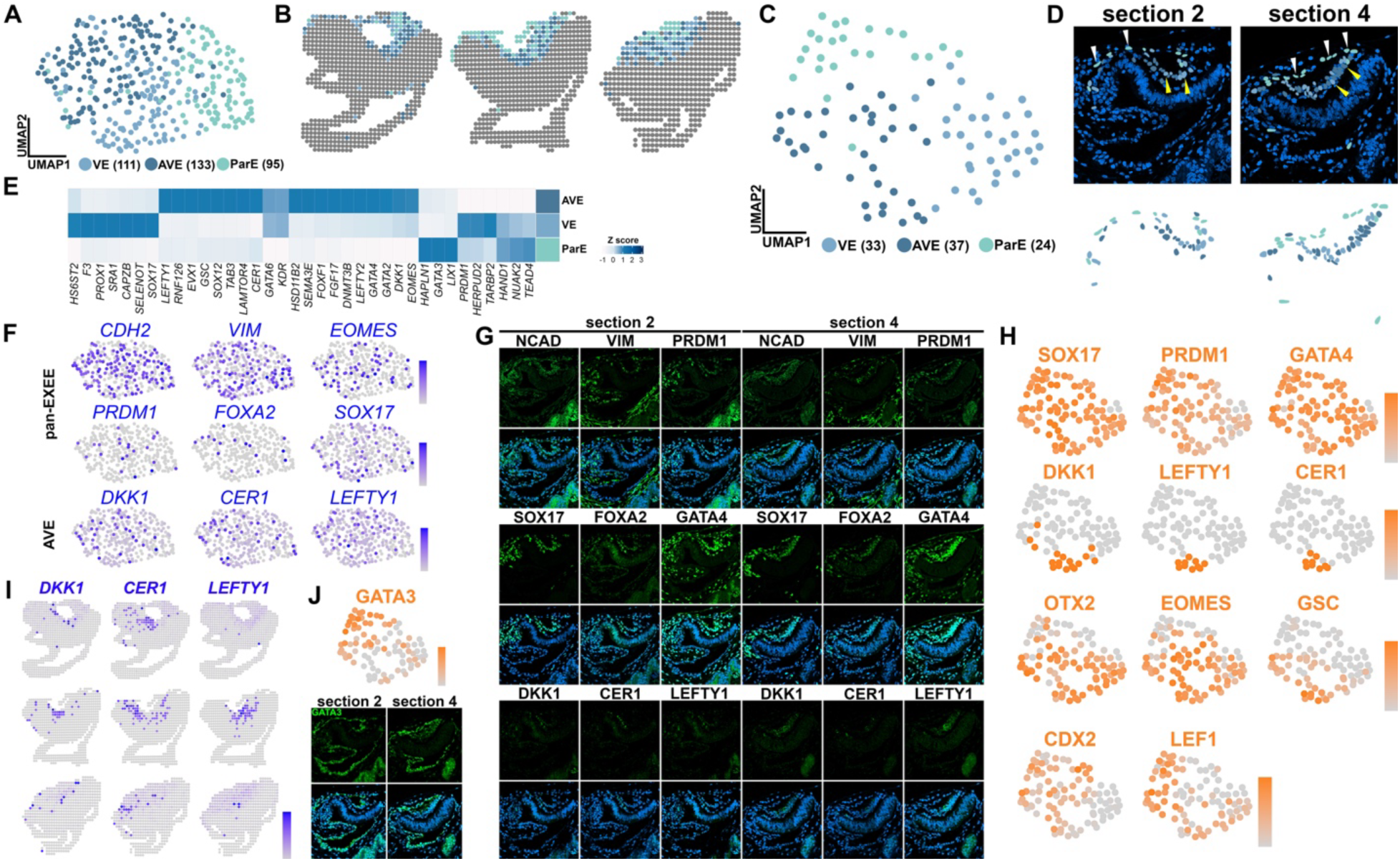
Sub-populations in the extra-embryonic endoderm. **A-D**) Transcriptomic and proteomic UMAP plots (A and C, respectively), displaying coordinates of each bin or cell with three identified populations: VE (light blue), AVE (blue), ParE (cyan), which are mapped spatially in (B, transcriptomic) and (D, proteomic). Numbers represent bins/cell per cluster. White arrowheads – ParE, yellow arrowheads – VE and AVE. **E)** A heatmap showing the normalized transcript expression values of selected endoderm lineage genes. **F)** Expression of selected lineage genes superimposed onto the transcriptomic UMAP plot (A). **G)** Confocal IF images of selected endoderm lineage marker from the proteomic dataset. **H)** Expression of indicated markers superimposed onto the proteomic UMAP plot in (C). **I)** Expression of *DKK1*, *CER1* and *LEFTY1*, AVE markers, superimposed onto the spatial plot. **J)** Top: Expression of GATA3 superimposed onto the proteomic UMAP plot in (C). Bottom: Confocal IF images of GATA3 in the sections 2 and 4.

Our understanding of ParE is currently limited. Morphologically, ParE cells (**Fig. 4D**, white arrowheads) are separated apart with flatter morphology compared to the VE cells (**Fig. 4D**, yellow arrowheads), and show the expression of organized TJP1, as well as PODXL and LAMA1 in apical and basal territories, respectively (**Fig. S8B**), which are key features of squamous epithelial cells [29, 68, 69]. Expression analyses show that the ParE and VE share the expression of several key genes (**Fig. 4F-H, S8C**, e.g., PRDM1, SOX17, GATA4). However, ParE cells do not show abundant expression of EOMES and GSC, nor secreted ligand inhibitors such as LEFTY1, DKK1 and CER1 (**Fig. 4E-I**). ParE cells also show nuclearly enriched CDX2 and LEF1 (**Fig. 4H**), known direct WNT targets [70–73], suggesting that WNT signaling is active in ParE cells, similar to mice [74]. Moreover, ParE enriched genes include *FGFR3*, *ID2*, and *LIFR* (**Fig. S8D**), known ParE genes in mice [75], as well as *GATA3*, a pioneer transcription factor [76–78] **Fig. 4E,J, S8E**), presenting GATA3 as a novel ParE marker. These results provide previously unrecognized molecular characteristics of primate ParE during early stages of gastrulation.

Expression analyses for selected amniotic markers in the transcriptomic and proteomic datasets show that most cells in the amnion territory display similar expression characteristics (**Fig. 5A-C**, also see **Fig. S9A**). Interestingly, in addition to the expression of known amnion markers (e.g., HAND1 and ISL1), TBX3 and TBX6, mesoderm markers [79–81] are expressed in the amnion (**Fig. 5B, S9B**). Notably, while PRDM14, a known marker of pluripotent cells [9, 82, 83], is rarely seen in the amnion tissue in the transcriptomic dataset, PRDM14 protein is abundant (**Fig. S9C**), suggesting that PRDM14 protein is highly stable. Moreover, cells that are positive for CLDN10, a marker for an amnion progenitor-like cell population that bridges the boundary between the amnion and epiblast [23], are seen anteriorly as well as posteriorly (arrowheads and dotted circles in **Fig. 5D**, also see **Fig. S9D,E** for nuclearly enriched pSMAD1/5 and the expression of *ID1*, *ID2*, *ID3* and *ID4* at the anterior and posterior tips). These results suggest that active amniogenesis may be present in both the anterior and posterior amnion-epiblast boundaries at CS6b.

**Figure 5.**
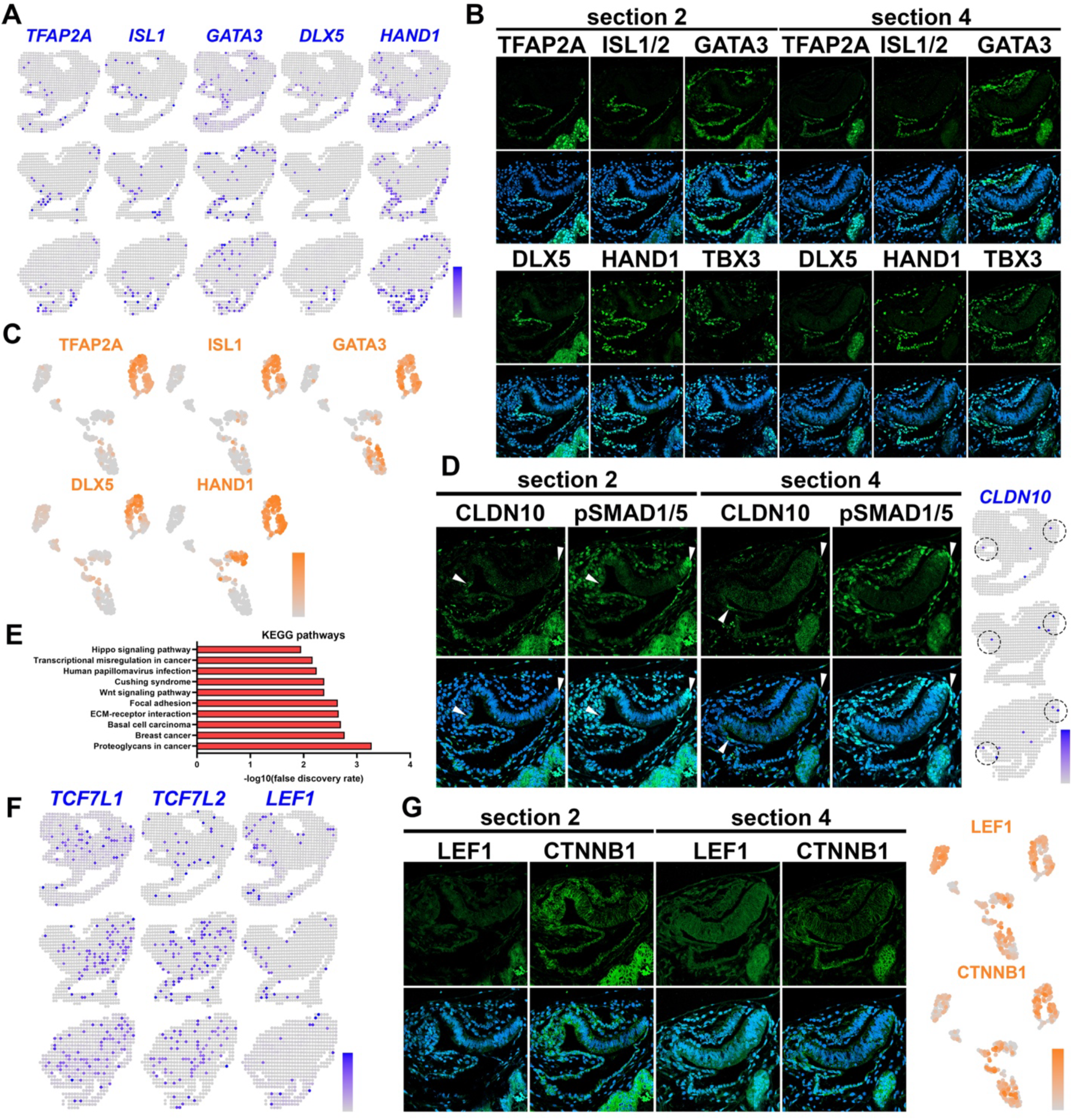
Transcriptomic and proteomic characterizations of the amnion. **A)** Expression of known amnion markers superimposed onto the spatial transcriptome plot. **B,C)** Confocal IF images of indicated amnion markers in the proteomic dataset (see (C) for the expression of these proteins in the UMAP space from Fig. 1E). **D)** Confocal IF images of the CS6b embryo stained for CLDN10 and pSMAD1/5. Arrowheads indicate CLDN10^+^ cells as well as spatially restricted nuclear pSMAD1/5 enriched cells (left). Expression of *CLDN10* superimposed onto the spatial transcriptome plot (right). Dotted circles indicate *CLDN10* enriched regions at the anterior and posterior epiblast-amnion boundaries. **E)** Top 10 KEGG pathways (ranked by FDR) based on all DEGs in the amnion (p<0.05) (**Table S10**). **F)** Expression of WNT-driven transcription factors superimposed onto the spatial transcriptome plot. **G)** Confocal IF images of immuno-stained sections of the CS6b embryo using LEF1 and CTNNB1 (top, spatial proteomic UMAP coordinates from Fig. 1E at the bottom).

**Figure 6.**
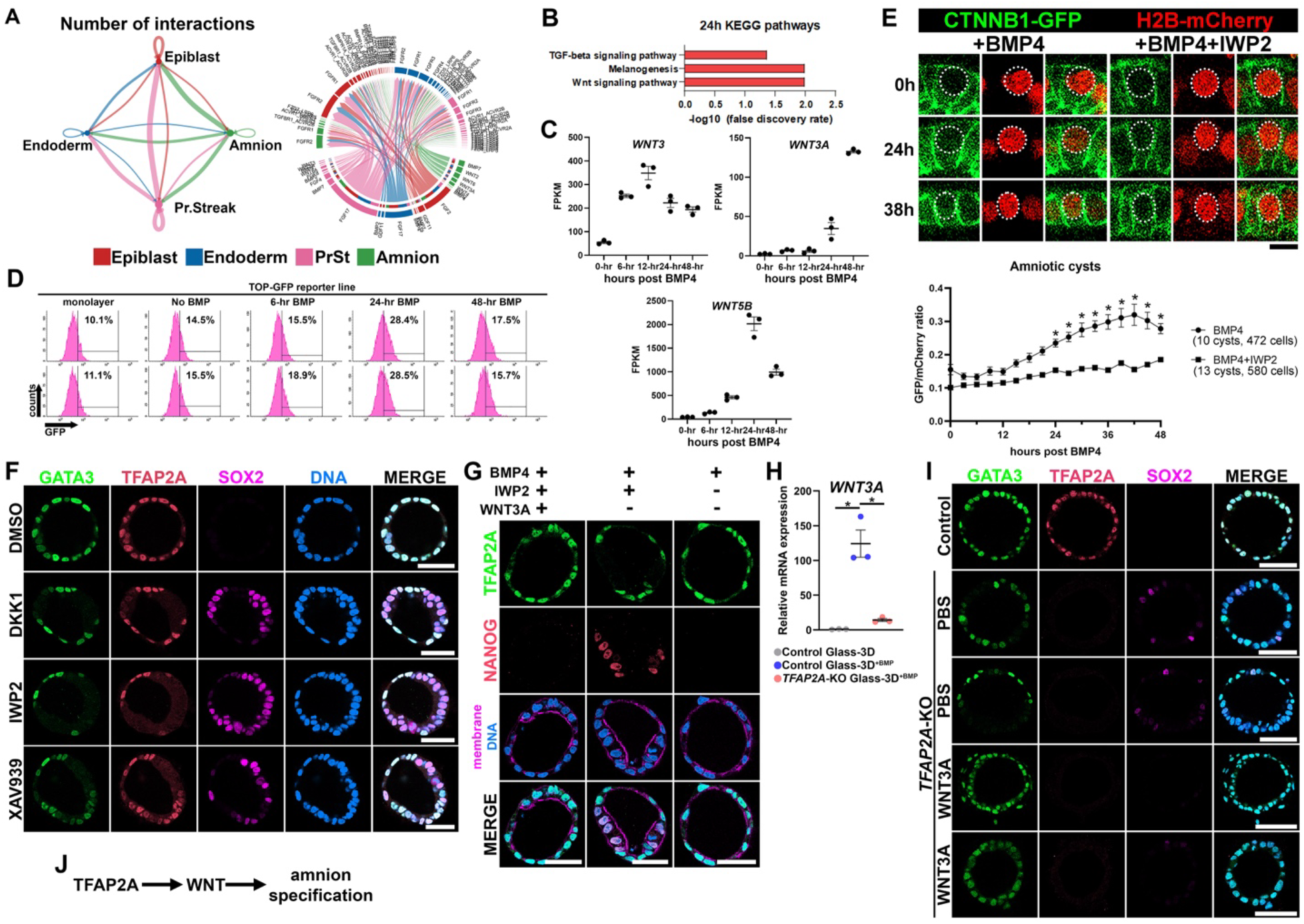
WNT signaling activity during amnion formation. **A)** Predicted receptor-ligand signaling networks among the four major cell types in the transcriptomic dataset. Thickness of the line indicates the signaling strength between cell lineages (left) or signaling strength of each receptor-ligand pair (right). **B)** Enriched pathways based on KEGG analysis of the DEGs (>2-fold change with p<0.05) at the 24-hr post-BMP4 timepoint in the Glass-3D^+BMP^ amniogenic model (based on our bulk-transcriptomic analysis in Sekulovski *et al.* [29]). **C)** Bulk-RNA sequencing time-course plots of normalized expression values for selected WNT ligands in developing Glass-3D^+BMP^ cysts. Time (in hr) represents timepoints after triggering amnion formation with recombinant BMP4. **D)** Original representative flow cytometry graphs of singly dissociated Glass-3D^+BMP4^ cells at indicated conditions, with cells gated for singlets and live cells. GFP^+^ cells were determined based on non-transgenic hPSC line. Percentages of GFP+ cells at each timepoint were calculated from the total of single and live cells. Representative plots are shown. **E)** Confocal time-lapse images (every 3 hours) of a representative CTNNB1-GFP knock-in/H2B-mCherry transgenic Glass-3D^+BMP^ amnion cysts with or without IWP2 (top). Scale bar 10µm. Quantitation is shown (bottom) for changes in nuclear fluorescent intensities (indicated by dotted circles) of GFP and mCherry signals (GFP/mCherry ratio) overtime, using ten control (472 total cells) Glass-3D^+BMP^ and thirteen IWP2-treated (580 total cells) Glass-3D^+BMP^ CTNNB1-GFP knock-in-H2B-mCherry cysts. Mean±SEM are shown, (One-Way ANOVA, post-hoc Dunnett) * p < 0.05. **F,G)** Confocal images of representative Glass3D^+BMP^ cysts at 48-hr post-BMP4 treatment, which are treated with indicated WNT signaling inhibitors (F), as well as with IWP2 and/or 200ng/mL hr-WNT3A (G) and stained with indicated markers. Scale bars 50µm. **H)** RNA expression of *WNT3A* in control pluripotent cysts, *TFAP2A*-KO and control Glass-3D^+BMP^ amniotic cysts, shown as mean±SEM, (Student’s t-test) * p <0.05. **I)** Optical sections of representative control and *TFAP2A*-KO Glass3D^+BMP^ cysts at 48-hr post-BMP4 treatment, treated with or without 200 ng/mL WNT3A. Scale bars 50µm. J) A schematic outlining the role of WNT signaling in amnion specification.

### WNT signaling-dependent amniotic fate progression during early gastrulation

Because the amnion has been shown to function as a signaling hub [3, 11, 12], we next examined signaling characteristics in the amnion territory. KEGG (Kyoto Encyclopedia of Genes and Genomes) analysis for all DEGs in the amnion reveals that WNT signaling is highly enriched (**Fig. 5E**, **Table S10**). Among the established WNT signaling transcription factors, *TCFL1*, *TCF7L2* and *LEF1* are abundantly expressed, and nuclearly enriched LEF1 and CTNNB1 are seen in the amnion (**Fig. 5F,G, S9F**).

Our analysis for receptor-ligand signaling activities based on CellChat ([84], **Fig. 6A, S10A-F**) shows that, in addition to identifying known signaling machineries associated with gastrulation (e.g., FGF, BMP, WNT) [85], this analysis predicts that signaling events are enriched among amniotic cells, as well as between the amnion and the epiblast or PrSt cells (**Fig. 6A**). Indeed, WNT signaling is highly abundant in the amnion cells (**Fig. S10D-F**, also previously shown in [8, 9]). These results suggest that WNT may be a key signaling program in early amniogenesis.

To experimentally examine the role of WNT signaling during amniogenesis we used a Glass-3D^+BMP^ amniogenic model [29], in which BMP4 ligand is added to pluripotent cell populations plated on a glass substrate (0-hr). Under these culture conditions, cysts with focally formed squamous amnion cells, surrounded by pluripotent cells, are formed by 20-30-hr post-BMP4 treatment, and fully squamous amnion cysts are observed by 48-hr [29]. Analysis of changes in bulk-transcriptomic characteristics in these developing Glass-3D^+BMP^ amnion cysts reveal an enrichment of highly expressed genes that are associated with WNT signaling at the 24-hr (intermediate) timepoint (**Fig. 6B**), including WNT ligands (e.g., *WNT3A*, *WNT5B*, **Fig. 6C**). Next, to investigate when WNT signaling is active, we performed time-course cytometry using a transgenic hPSC line that expresses GFP under the control of a 6x*TCF/LEF* promoter [86], as well as time-lapse imaging using an hPSC line with mEGFP knocked into the *CTNNB1* (also known as β-catenin) loci (Allen Institute, AICS-0058)); this line also expresses mCherry-H2B [87] to label nuclei. Flow cytometry analysis of the 6x*TCF/LEF*;*GFP* line shows that GFP^+^ cells are most enriched at the 24-hr timepoint (**Fig. 6D, S11A,B**). Time-lapse imaging followed by quantitation for the nuclear GFP/mCherry fluorescent signal intensity ratio shows a significant increase in the level of nuclear mEGFP-CTNNB1 starting at 24-hr, which then peaks at 42-hr; this increase is not seen when cysts are treated with IWP2 starting at 0-hr (when BMP is also added), a small molecule inhibitor of Porcupine, an intracellular enzyme critical for WNT ligand secretion and function [88] (**Fig. 6E**). Therefore, WNT signaling is active by the intermediate (20-30-hr) timepoint during amniogenesis in the Glass-3D^+BMP^ system.

Next, developing Glass-3D^+BMP^ cysts were treated with recombinant DKK1, a secreted WNT ligand inhibitor, or IWP2 starting at 0-hr and then harvested at 48-hr. Strikingly, while DMSO-treated controls show uniformly squamous amniotic cysts at 48-hr; DKK1– or IWP2- treated cysts contain both squamous (TFAP2A^+^) and columnar (SOX2^+^) cell populations (**Fig. 6F**), an organization similar to that of intermediate timepoints in Glass-3D^+BMP^. Cysts treated with XAV939, a small molecule inhibitor of canonical WNT signaling which inhibits tankyrase activity [89], also show this intermediate amnion cyst organization (**Fig. 6F**). In contrast, IWP2-treated cysts that are also treated with recombinant WNT3A express TFAP2A, but not NANOG, in all cells and are fully squamous in nature (**Fig. 6G**), indicating that canonical WNT signaling is critical for amniogenesis.

Importantly, this intermediate phenotype seen after WNT signaling inhibition is similar to that seen in Glass-3D^+BMP^ cysts lacking TFAP2A [29]. Indeed, quantitative PCR analysis reveals a significant reduction in the expression of *WNT3A* in *TFAP2A*-KO Glass-3D^+BMP^ cysts compared to controls (**Fig. 6H**). Moreover, although sporadic GATA3^+^ mesenchyme-like cells are seen in the periphery of the developing cysts, treatment of *TFAP2A*-KO cysts with recombinant WNT3A rescues the intermediate cyst phenotype, resulting in fully squamous cysts (**Fig. 6I**). Together, these results demonstrate that TFAP2A-driven canonical WNT signaling is critical for maintaining amniogenesis (**Fig. 6J)**.

## Discussion

In summary, we developed a spatially-resolved multiomic resource of the early primate gastrula with significantly greater spatial and molecular detail than previous studies [9, 19, 20], which allows identification of previously unrecognized cell types and developmental features in several early embryonic lineages (see **Fig.7** for all identified cell types that are superimposed onto the spatial maps). The fact that Vimentin transcript and protein levels differ dramatically in a tissue-specific manner reveals the presence of a tissue-specific translational mechanism, illuminating such mechanism as a potential driver of lineage progression and presenting a need to globally identify pre– and post-translational levels of gene products. Moreover, the identification of transcriptomic and proteomic similarities between PGC and endoderm populations (nDE, VE and AVE) opens an avenue for future investigations into mechanisms by which these cell populations integrate during early embryogenesis. Finally, while previous studies using stem cell models have not conclusively supported or rejected a role for WNT signaling during amniogenesis [9, 90–92], our Glass-3D^+BMP^ platform robustly exposes an amniotic requirement for WNT signaling, perhaps due to its single lineage nature. Broadly, this is a robust spatial multiomic pipeline that can be utilized to globally investigate a variety of aspects of embryonic development and disease.

**Figure 7.**
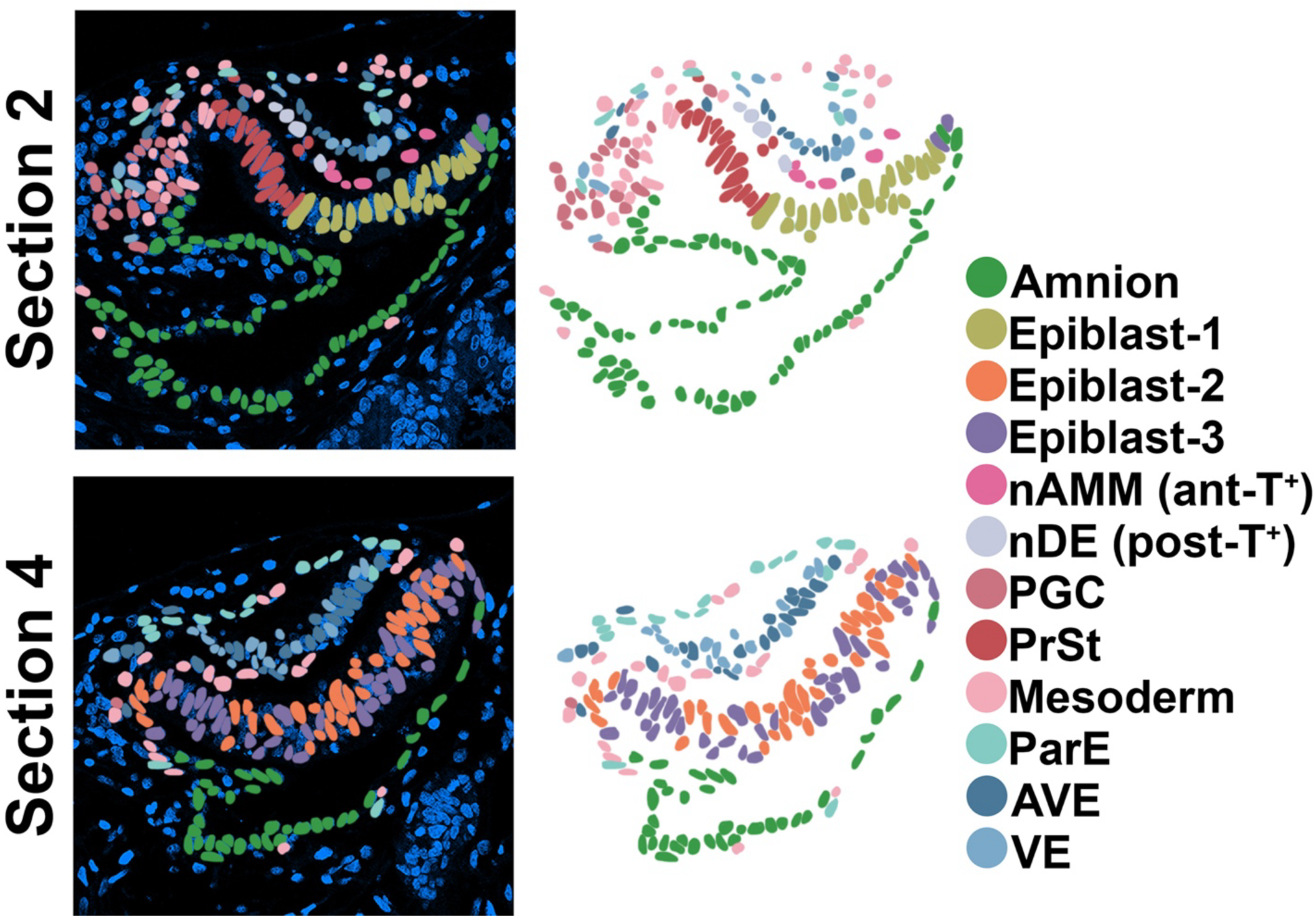
Cell lineages in an early-mid primate gastrula. Newly identified lineages are annotated in the sections 2 and 4 with indicated colors.

## Acknowledgments

We thank the Wisconsin National Primate Research Center (WNPRC) Veterinary, Scientific Protocol Implementation, Pathology and Colony Services staff for providing animal care, and assisting in procedures including breeding, pregnancy monitoring, and sample collection. A special thanks to Drs. Heather Simmons and Puja Basu for collecting the macaque specimens. The contents of this manuscript are solely the responsibility of the authors and do not represent the official views of the NIH. We thank Drs. Deborah Gumucio and William Shawlot for insightful comments to the manuscript, as well as the Roy J. Carver Biotechnology Center at the University of Illinois at Urbana-Champaign for sequence services. This work was supported by NIH grants R01-HD098231 (K.T.), P51 OD011106 (to the WNPRC) as well as by MCW CBNA Start-up funds, Advancing a Healthier Wisconsin (AHW) Endowment (16003-5520766, N.S.) and the Lalor Foundation Postdoctoral Fellowship (N.S.).

## Materials and methods

### Cynomolgus macaque

#### Animals

The female and male cynomolgus macaques were housed and cared for at the Wisconsin National Primate Research Center (WNPRC). All procedures were performed in accordance with the NIH Guide for the Care and Use of Laboratory Animals and under approval of the Institutional Animal Care and Use Committee (protocol g005061) at the University of Wisconsin-Madison.

#### Animal breeding and pregnancy detection

The female was housed with a compatible male between days 6-17 post-onset of menses and monitored for breeding. On days 13, 15 and 17 post-onset of menses, 2 mL of blood was withdrawn from the saphenous vein to isolate serum. Serum estradiol and progresterone levels were quantified using a cobas e411 analyzer as previously described [29]; luteal levels of estrogen and progesterone were observed at all timepoints suggesting that ovulation occurred prior to day 13. Two transabdominal ultrasounds were performed to detect pregnancy and estimate gestational age based on gestational sac measurements [93].

#### Terminal perfusion uterine collection, paraffin embedding and sectioning

The pregnant female was sedated with intramuscular ketamine (>15 mg/kg) followed by IV sodium pentobarbital (>35 mg/kg) and then perfused with 4% paraformaldehyde (PFA) via the left ventricle. The entire uterus and cervix were removed. The serosa and superficial myometrium were scored for better fixative penetration and to denote dorsal and ventral orientation. The tissues were fixed in 4% PFA. The uterus was serially sectioned from the fundus to the internal cervical os into 4 mm slices using a dissection box after 24 hrs in fixative. The tissues within cassettes were placed in fixative for a total of 72-hrs in fixative. Cassettes with tissue sections were transferred into 70% ethanol, routinely processed in an automated tissue processor and embedded in paraffin for histological analyses (5µm sections).

### Visium HD spatial transcriptome

5µm uterine sections were placed on a histology slide. Tissue sections were subjected to deparaffinization, nuclear staining (Hoechst 33342), and imaging as described in the manufacturer’s protocol (Visium CytAssist Spatial Gene Expression for FFPE – Deparaffinization, Decrosslinking, Immunofluorescence Staining & Imaging Handbook (CG000584, 10x Genomics)). Probe hybridization, probe ligation, Visium HD slide preparation, probe release, extension, library construction, and sequencing was performed as described in the manufacturer’s protocol (Visium HD Spatial Gene Expression Reagent Kits User Guide (CG000685, 10x Genomics)). Briefly, probe hybridization was performed using human whole transcriptome probe panel, consisting of ∼3 specific probes per target gene (PN-1000466, 10x Genomics). After hybridization, the Probe Ligation Enzyme (PN-2000425, 10x Genomics) was added to generate probe-RNA ligation products, which was followed by probe release (from the histology slide) and capture (onto the Visium HD slide) using Visium CytAssist (Ser.# CAVG10689, Firmware 2.4, 10x genomics). The ligation products were then extended and eluted, followed by amplification and indexing using sample index PCR. The final libraries were cleaned up by SPRIselect. Sequencing was performed on an Illumina NovaSeq X (10B paired-reads 300-cycle) to obtain paired-end reads. Space Ranger version 3.0 was used for analysis. In addition to the native 2µm tile size, Space Ranger outputs Visium HD data binned to 8µm and 16µm resolution. Downstream analyses were performed using the 8µm-binned data.

### Spatial proteomics based on FFPE-4i

#### Paraffin-embedded cynomolgus macaque embryo section standard immunostaining

Fixed uterine tissue were cut in 5 µm thick cross-sections, mounted on slides, deparaffinized in xylene, and rehydrated in an ethanol series. Antigen retrieval was performed by boiling in citrate buffer. Sections were blocked using 4% donkey serum in PBS at RT for at least 3 hours. Immunolocalization was performed using commercially available primary antibodies, incubated overnight at 4 °C in 4% serum, followed by incubation with secondary antibodies tagged with a fluorescent dye (fluorophores excitation = 488, 555, and 647 nm), which are then counterstained with DAPI. Negative controls were performed in which the primary antibody was substituted with the same concentration of normal IgG of the appropriate isotype. Cover-slipped samples were imaged using a Zeiss LSM980 microscope. Antibodies for IF staining are found in **Table S11**.

#### Multiplex immunostaining, imaging and mounting/unmounting of cynomolgus macaque paraffin-embedded sections (FFPE-4i)

We adopted a multiplex immunostaining protocol previously developed by Gut *et al.* [22] for FFPE samples that require mounting [23]. In summary, following IF staining imaging (tile imaging, using a Zeiss LSM980 microscope, followed by image alignment and processing in Photoshop), glass slides with coverslip-mounted samples were immersed in water for 5 min to dilute mounting medium. It is recommended to unmount within 12 h of mounting (and imaging) before mounting medium dries and/or hardens. After unmounting, slides were washed six times with water (3 min each). Slides were then incubated in Denaturing Buffer with gentle rock (∼ 100 rpm) for 10 min (three times), followed by incubation in Stripping Blocking Solution with gentle rocking (∼ 100 rpm) for 1 h. Next, slides were washed 3 times (5 min each) in PBS; this step was followed by standard immunostaining (described previous section): primary antibody incubation, washing, secondary antibody incubation/counter staining, washing and mounting. Samples were immediately imaged. All steps were performed at room temperature. Denaturing Buffer—0.5M L-Glycine (Sigma Aldrich, cat: G8898), 3M Urea, 3M Guanidine hydrochloride (Sigma Aldrich, cat: G4505), and 70 mM TCEP-HCl (TCEP) (Vector, cat: QBD-10014) in ddH_2_0. Adjust to pH 2.5, and store at 4 °C for up to 2–3 months. This solution can be reused several times. Stripping Blocking Solution—4% Donkey Serum (Millipore, cat: S30) and 150 mM Maleimide (Sigma Aldrich, cat: 129,585) in phosphate buffered saline (PBS). Store at 4 °C, and needs to be used within 1 week. This is a single use solution.

#### Quantification of FFPE-4i

Each cell within the two sections of the embryo was labeled (section 2, total of 216 cells; section 4, total of 191 cells) (**Fig. 1E,N**). All IF images from each section were aligned, and the intensity of each nuclear signal was quantified using ImageJ (NIH); the quantitation was done in blind.

The intensity matrix was used for unsupervised clustering using Seurat. Data was normalized using CLR, followed by FindVariableFeatures (45 proteins), and ScaleData. Cell clusters were identified using FindNeighbors (1:10 dimensions) and FindClusters functions based on a shared nearest neighbor modularity optimization clustering algorithm (resolution set as 0.4).

### hESC lines

Human embryonic stem cell line H9 (WA09, P30, P48, WiCell; NIH registration number: 0062) and induced pluripotent stem cell line AICS-0058-067 (mEGFP-CTNNB1 knock-in) were used in this study. All protocols for the use of hPSC lines were approved by the Human Stem Cell Research Oversight Committee at the Medical College of Wisconsin. All hPSC lines were maintained in a feeder-free system for at least 20 passages and authenticated as karyotypically normal at the indicated passage number. Karyotype analysis was performed at Cell Line Genetics. All hPSC lines tested negative for mycoplasma contamination (LookOut Mycoplasma PCR Detection Kit, Sigma-Aldrich). In summary, hESC were maintained in a feeder-free culture system with mTeSR1 medium, or with 50%/50% mix of mTeSR1 and mTeSR plus (STEMCELL Technologies). hESC were cultured on 1% (v/v) extracellular matrix (ECM, e.g., Geltrex (Thermo Fisher Scientific), Cultrex SCQ (Bio-techne)) coated 6 well plates (Nunc). Cells were passaged as small clumps every 4 to 5 days with Dispase (Gibco). All cells were cultured at 37°C with 5% CO_2_. Media was changed every day. hESC were visually checked every day to ensure the absence of spontaneously differentiated mesenchyme-like cells in the culture. Minor differentiated cells were scratched off the plate under a dissecting scope once identified. The quality of all hESC lines was periodically examined by immunostaining for pluripotency markers and successful differentiation to three germ layer cells. Similar methods were previously used in [29, 94].

#### piggyBac-based transgenic edited lines

PB constructs (3 μg, **Table S11**) and pCAG-ePBase [95] (1 μg) were co-transfected into H9 hESC or AICS-0058-067 cells (70,000 cells cm^−2^) using GeneJammer transfection reagent (Agilent Technologies). To enrich successfully transfected cells, drug selection (puromycin, 2 µg/mL for 4 days) was performed 48 to 72 hours after transfection. Cells stably expressing each construct maintained the expression of pluripotency markers.

### Glass-3D^+BMP^ amnion cyst assays

#### Amnion cyst formation

This assay has been previously described [29]. Briefly, singly dissociated cells were prepared using Accutase (Sigma-Aldrich) and were plated on 1% ECM-coated coverslips at 25,000 cells/cm^2^ in mTeSR1 or in 50%/50% mTeSR1/mTeSR plus mix medium in the presence of 10µM Y-27632 (STEMCELL Technologies). After 24 hr (day 1), cells were then incubated in media containing 2% ECM overlay, without Y-27632. At 48 hr post plating (day 2) cells were incubated in media containing 1% ECM with 20nM BMP4, followed by daily media changes with 1% Geltrex, without BMP4. Small molecule treatment was performed at indicated concentrations (**Table S11**) from day 2 to day 4.

#### Immunostaining

Glass-3D^+BMP^ cysts grown on the coverslip were rinsed with PBS (Gibco) twice, fixed with 4% paraformaldehyde (Sigma) for 60 min, then rinsed with PBS three times, and permeabilized with 0.1% SDS (Sigma, in 1x PBS) solution for 60 min. The samples were blocked in 4% heat-inactivated goat serum (Gibco) or 4% normal donkey serum (Gibco) in PBS overnight at 4°C. The samples were incubated with primary antibody solution prepared in blocking solution at 4°C for 48-hr, washed three times with PBS (30 min each), and incubated in blocking solution with goat or donkey raised Alexa Fluor-conjugated secondary antibodies (Thermo Fisher), at room temperature for 24 hours. Counter staining was performed using Hoechst 33342 (nucleus, Thermo Fisher Scientific). All samples were mounted on slides using 90% glycerol (in 1x PBS). When mounting hPSC-cysts samples, modeling clay was used as a spacer between coverslip and slide to preserve lumenal cyst morphology. Samples were acquired using a Zeiss LS980 laser scanning confocal microscope. Images were generated using Zen, FIJI (NIH) and Photoshop (Adobe).

#### RNA isolation and qRT-PCR analysis

Total RNA extraction was performed using TRIzol (Thermo Fisher). The quantity and quality of total RNA was determined by spectrometry and denaturing agarose gel electrophoresis, respectively. The cDNA was synthesized from total RNA (1 μg) using High-Capacity cDNA Reverse Transcription Kit (Applied Biosystems). Quantitative real-time RT-PCR (qPCR) analysis of mRNA expression was performed using a CFX96 Real-Time PCR Detection System (Bio-Rad) with Power Up SYBR Green Master Mix (Applied Biosystems). Primers (**Table S11**) were designed using NCBI primer design tool to span at least one exon/intron junction. Data were normalized against GAPDH and are shown as the average fold increase ± standard error of the mean (SEM) with relative expression being calculated using the 2-ΔΔCT method as described previously [96]. After amplification, the specificity of the PCR was determined by both melt-curve analysis and gel electrophoresis to verify that only a single product of the correct size was present.

### Bioinformatics

Analysis of the Visium HD dataset was performed using the R package Seurat (v.4.5.0, [37]). For quality control, bins with fewer than 180 detected genes were excluded. Expression values of 8µm bins were then log-normalized with a scaling factor 10,000. Samples were then integrated using an anchor-based integration approach implemented by FindIntegrationAnchors function (dimensions 1-10) followed by IntegrateData function. The integrated data was subsequently centered and scaled across genes. Principal component analysis (PCA) was performed, and the top 50 principal components were used for two-dimensional UMAP embedding of 1,998 bins (744, 690, and 564 bins in sections 1, 3, and 5, respectively). Clusters of 8µm bin clusters were identified using the FindNeighbors and FindClusters functions based on a shared nearest neighbor modularity optimization clustering algorithm (resolution set as 0.5). Cluster marker genes were identified using the FindAllMarkers function (Wilcoxon rank sum test) with adjusted p-values less than 0.05.

#### SCENIC analysis

Gene regulatory network analysis was performed using SCENIC [97] implemented by python package pySCENIC, using a three-step approach based on human cisTarget resources. Firstly, the gene expression matrix from 8µm bins was converted to a Loom file format using the loompy (3.0.8) python library. Next, ‘pyscenic grn’, ‘pyscenic ctx’ and ‘pyscenic aucell’ functions were used to infer the gene regulatory network, estimate regulon activity, and identify gene regulatory networks associated with major transcription factors and their target genes. Regulon activity scores were computed for individual bins and summarized at the cluster level. Regulon specificity scores (RSS) were calculated for each cluster to quantify cluster-specific regulatory activity. Cell type-specific regulons were identified by selecting top-ranked regulons based on the RSS values, and visualized using RSS plots.

#### Hotspot analysis

Spatial coordinates of each spot and the regulon activity matrix from SCENIC analysis were used to construct a spot similarity graph by create_knn_graph from python library hotspotsc [34] to capture local neighborhood relationships among spatial bins. Subsequently, features that exhibit significantly non-random expression patterns in the similarity graph were selected based on significant local autocorrelation (p-values < 0.05). Pairwise associations among these spatially variable features were then conducted to construct a Z-score matrix, analogous to a correlation matrix, which was subsequently clustered to identify coherent gene/regulon modules based on shared local spatial correlation. Module-level activity scores were computed on a per-spot basis using the hs.calculate_module_scores function, and the resulting module scores were integrated into our dataset. Finally spatial distributions of regulon module activities were visualized for each tissue slice using the SpatialFeaturePlot function according to spatial coordinates.

#### Pseudotime analysis

Pseudotemporal lineage trajectories were inferred using the R package Monocle3 [32]. A principal graph representing cellular state transitions was learned using the learn_graph function with parameters minimal_branch_len = 15 and geodesic_distance_ratio = 1.5. The trajectory root was defined by selecting an appropriate root node corresponding to the inferred cell origin. Pseudotime values were then assigned to individual cells by ordering cells along the learned trajectory in the UMAP embedding, using the geodesic distance from the selected root node. Gene expression dynamics along the inferred trajectories were visualized using the plot_genes_in_pseudotime function.

#### CellChat analysis

After cluster annotation, the R package CellChat [84] was applied to explore cell-cell communication network. The normalized 8µm bin gene expression matrix and annotated spatial cell-type information served as the input for CellChat analysis. Communication probabilities between the cell types were computed using a trimming threshold of trim = 0.02 to reduce the influence of lowly expressed interactions. CellChatDB.human database was used to identify statistically significant receptor-ligand interactions (p < 0.05) and to characterize signaling pathways involved in intercellular communication, including the WNT, BMP and FGF pathways. Finally, gene expression data was integrated with cell-cell communication probabilites and interaction strengths to visualize predicted spatial communication patterns.

#### STdeconvolve analysis

We used a reference-free deconvolution tool STdeconvolve [24] to assess whether our Visium HD data (8 µm bins) contained potential cell-type mixtures within individual bins. We fit latent Dirichlet allocation (LDA) models across a range of topic numbers (K = 2–10) and evaluated model diagnostics, including perplexity and the number of inferred topics with non-trivial contributions (mean proportion > 5%). In our dataset, perplexity increased monotonically with increasing K, while the number of topics exceeding the 5% mean proportion threshold remained near zero across all tested values. Together, these diagnostic patterns indicate that increasing model complexity does not improve explanatory power and suggest that most bins behave as near single-cell–level expression profiles rather than mixed-cell profiles.

#### Pairwise spatial neighborhood analysis

To assess whether specific pairs of clusters exhibited non-random joint spatial organization within each spatial field of view, we performed a permutation-based spatial neighborhood analysis. Spatial neighborhoods were defined using a k-nearest-neighbor (kNN) graph constructed from two-dimensional spatial coordinates. For a given pair of clusters (e.g., Epiblast-2 and Epiblast-3), we defined a local joint spatial organization statistic by computing, for bins assigned to one cluster, the fraction of their kNNs that were assigned to the other cluster. To generate a null distribution of the statistic, cluster labels for the selected cluster pair were randomly permuted within each field of view while preserving spatial coordinates and cluster sizes, and the statistic was recomputed under the null across 5,000 independent permutations. Statistical significance was assessed by comparing the observed statistic to the permutation-derived null distribution, allowing detection of non-random local spatial mixing or segregation between the two clusters.

#### RNA and protein integration

To integrate the transcriptomic (Visium, 10x Genomics) and proteomic (FFPE-4i) datasets, the spatial transcriptomics object was first subset to retain genes overlapping with the protein targets measured in the proteomic dataset. Transcriptomic data were normalized using centered log-ratio (CLR) normalization to ensure comparability with the CLR-normalized proteomic data. Two datasets were then merged, and aligned using an anchor-based integration framework by the IntegrateData function from the R package Seurat [37]. Principal component analysis (PCA) was performed on the integrated data, and the top 12 principal components were used for two-dimensional UMAP embedding. Cell neighborhood relationships were identified using the FindNeighbors function (dimensions 1-4), followed by clustering analysis with FindClusters based on a shared nearest-neighbor (SNN) modularity optimization algorithm.

#### Statistical analysis

Graphs were generated using Prism 7 (GraphPad Software) or R, and statistical analyses were performed using Prism 7. We used Student’s t-test or ANOVA with post hoc Dunnet’s multiple comparisons test, as indicated in figure legends. In **Fig. S1Q**, eight independent measurements were taken for each sample. In **Fig. 6E**, 10 and 13 independent cysts were imaged, with a total number of 472 and 580 imaged cells. In **Fig. 6H**, RNA was collected from 3 independent samples. In **Fig. S7G**, nuclear immunofluorescence intensity was measured in all indicated cells. Statistical significance was indicated with * p-value < 0.05.

## Supplementary table legends

**Table S1. Spatial transcriptomics – 10X Visium HD human probes for the use with cynomolgus macaque (*Macaca fascicularis*)**. **A-D**) List of probes that detect *Macaca fascicularis* genes (provided by the manufacturer). List of probes that detect a single gene (A) or no gene (B). Probes that are predicted to display multiple predicted hits are shown in (C). Mixed multi-probe (D) contains multiple probes for the same human gene, but not all of those probes are predicted to detect the gene in *Macaca fascicularis*.

**Table S2. Spatial proteomics – fluorescent intensity matrix.** IF-stained FFPE-4i samples were imaged and were quantitated for fluorescent intensity. Nuclear intensity quantiation was performed for all markers except for membrane (CLDN10, CDH1, CDH2), secreted (LEFTY1, DKK1, CER1) and cytosolic (VIM) proteins, in which we quantitated binary (1 (positive); 0 (negative)) for each cell.

**Table S3. Spatial transcriptomics – 8μm bin matrix.** Normalized counts per bin, used for spatial transcriptomics data.

**Table S4. Spatial transcriptomics – differentially expressed genes in the four major cell populations. A)** Epiblast, **B)** Amnion, **C)** Primitive Streak and **D)** Endoderm.

**Table S5. Spatial transcriptomics – AUCell matrix for 322 enriched regulatory networks for all 8μm bins, based on SCENIC**

**Table S6. Spatial transcriptomics – List of enriched gene regulatory networks in each of the 8 modules, based on Hotspot. A-H)** Modules 1 through 8, respectively. RSS score for each gene regulatory network is shown for each bin.

**Table S7. Spatial transcriptomics – Gene set enrichment analysis (GSEA) of module-enriched gene regulatory networks. A-H)** Modules 1 through 8, respectively.

**Table S8. Spatial transcriptomics – differentially expressed genes in Epiblast-1, Epiblast-2, Epiblast-3, Primitive Streak (PrSt), Mesoderm, nascent definitive endoderm and PGC (nDE/PGC) clusters (labeled A-F, respectively)**

**Table S9. Spatial transcriptomics – Differentially expressed genes in the endoderm sub-clusters (ParE, A; VE, B; AVE, C)**

**Table S10. Spatial transcriptomics – GSEA of DEGs enriched in the amnion cluster**

**Table S11. Small molecules (A), antibodies (B,C), plasmids (D), primers (E) and cell lines (F) used in this study**

## Figures and figure legends

**Figure S1.**
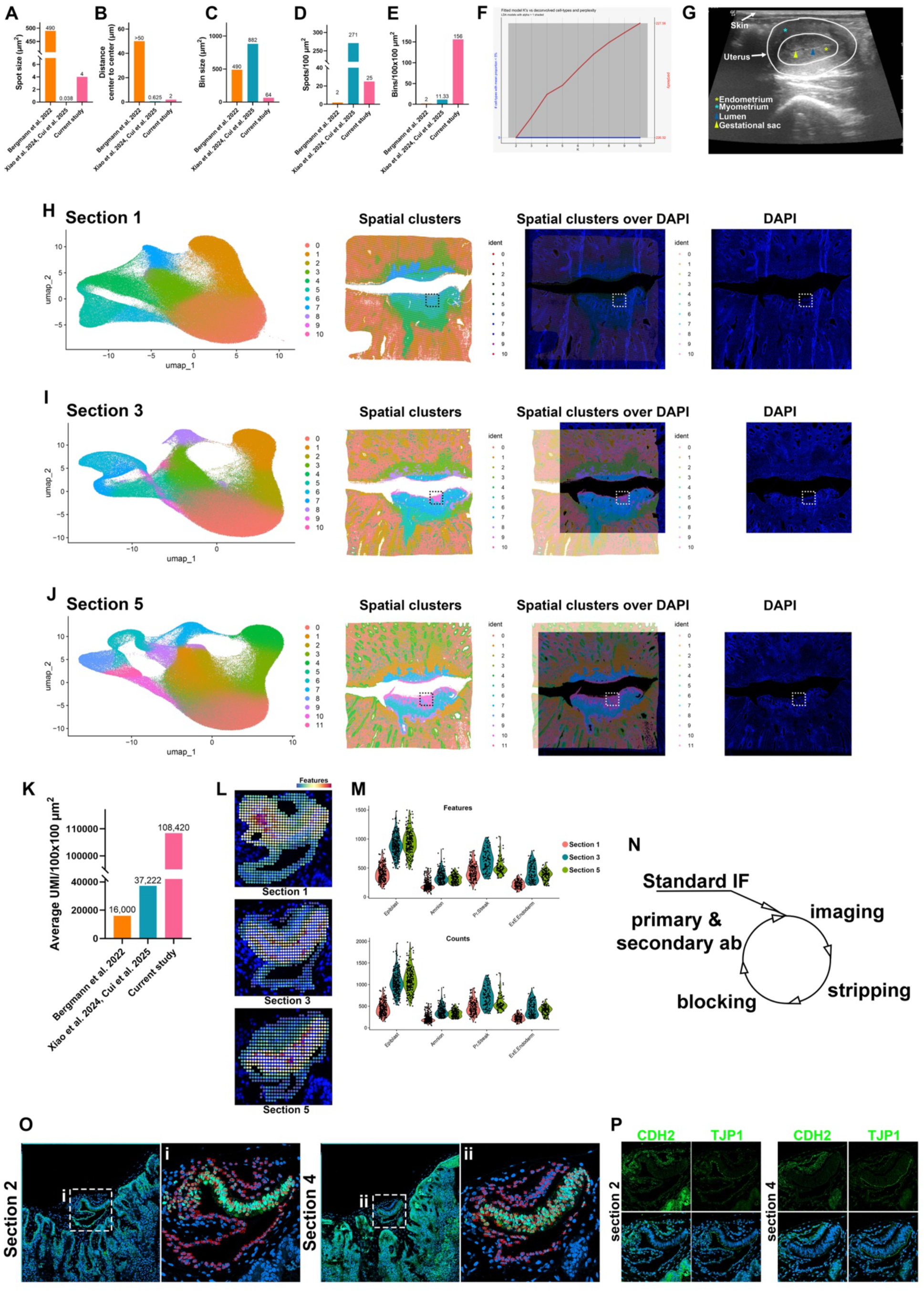
Additional information for the transcriptomic and proteomic pipelines. **A-E)** Comparisons of spatial transcriptomic resolution across indicated studies (including this study) that investigated early primate development (A, spot size (smallest unit in spatial transcriptomic analysis); B, center-to-center distance of neighboring spots; C, bin size; D, spot density; E, bin density). **F)** Perplexicity analysis of the deconvoluted bins using STdeconvolve. **G)** An abdominal ultrasound image showing the uterus with a gestational sac. **H-J)** UMAP plots (1^st^ column) showing the coordinates of all bins in the entire scanned area (2^nd^ column) with cluster identities. The scanned area was stained for DNA (blue, shown in the 3^rd^ and 4^th^ columns). Images in the 2^nd^ column were superimposed onto the DNA-stained images (3^rd^ column). Dotted boxed area indicate the regions that are analyzed in detail in this study. **K)** A bar graph showing the average UMI per 10^4^
μm^2^. **L,M)** Number of unique genes (Features, top) and transcripts (Counts, bottom) for each bin is plotted in the spatial transcriptomics plot (L) as well as per cell lineage (M). **N)** A schematic outlining the FFPE-4i pipeline. **O)** Confocal IF images of the entire scanned region for the FFPE-4i analysis (sections 2 and 4, stained with CDH1 and DNA). Insets (i,ii) indicate regions that are used for the spatial proteomic analysis (stained with OCT4 and DNA); numbers provide cell identity during quantitation. **P)** Confocal IF images of the sections 2 and 4, stained with indicated polarity markers.

**Figure S2.**
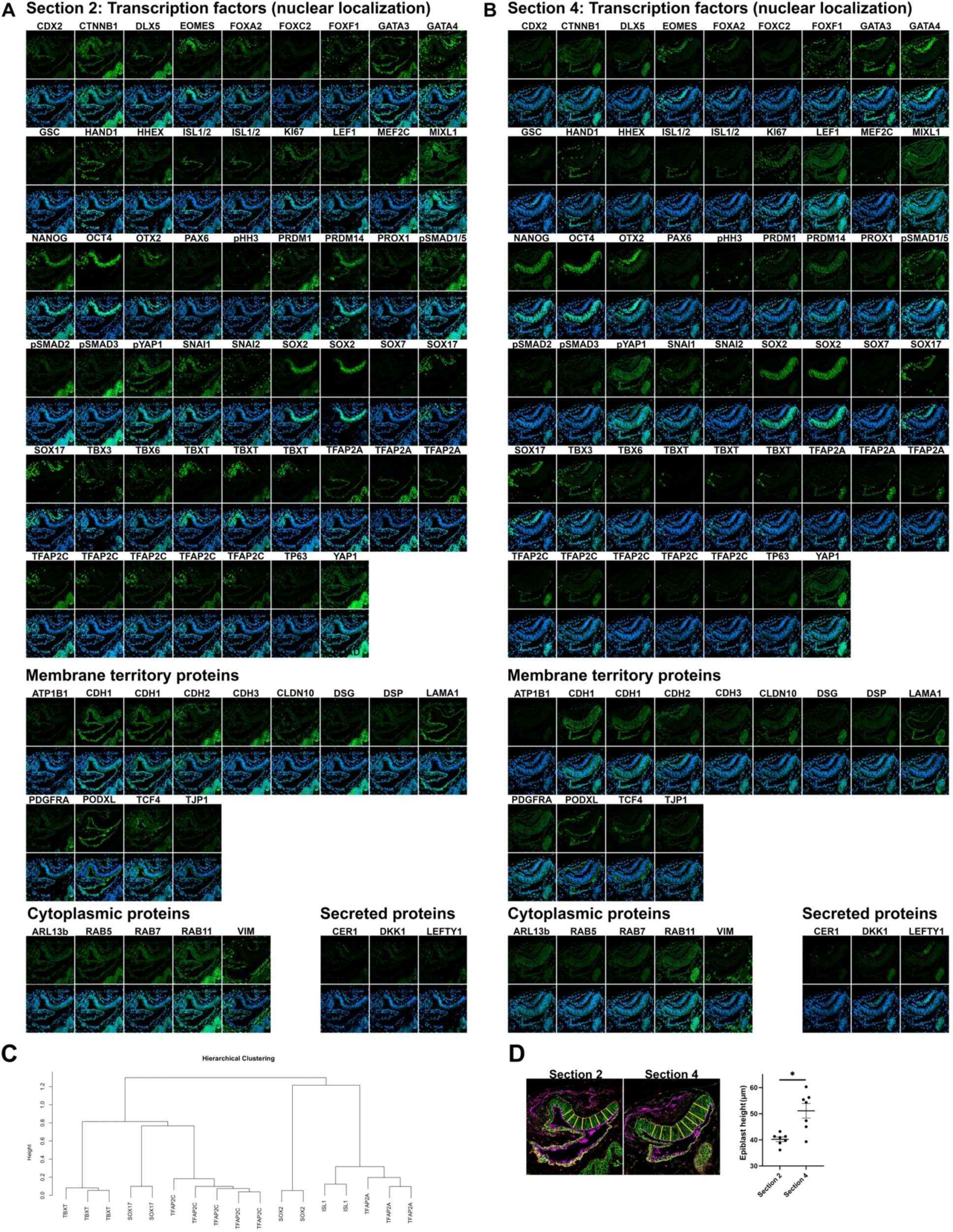
Additional information for the spatial proteomics. **A,B**) Confocal IF images for all tested proteins in the spatial proteomic dataset. **C)** Replicate marker reproducibility analysis based on hierarchical clustering. A hierarchical clustering plot showing that the markers that were used in more than one FFPE-4i cycle display similar expression intensity/pattern. **D)** The pipeline to measure the epiblast cell height (left). A scatter plot showing the length of the eight yellow lines (epiblast height) in the section 2 and section 4. Mean±SEM are shown, (Student’s t-test) * p < 0.05.

**Figure S3.**
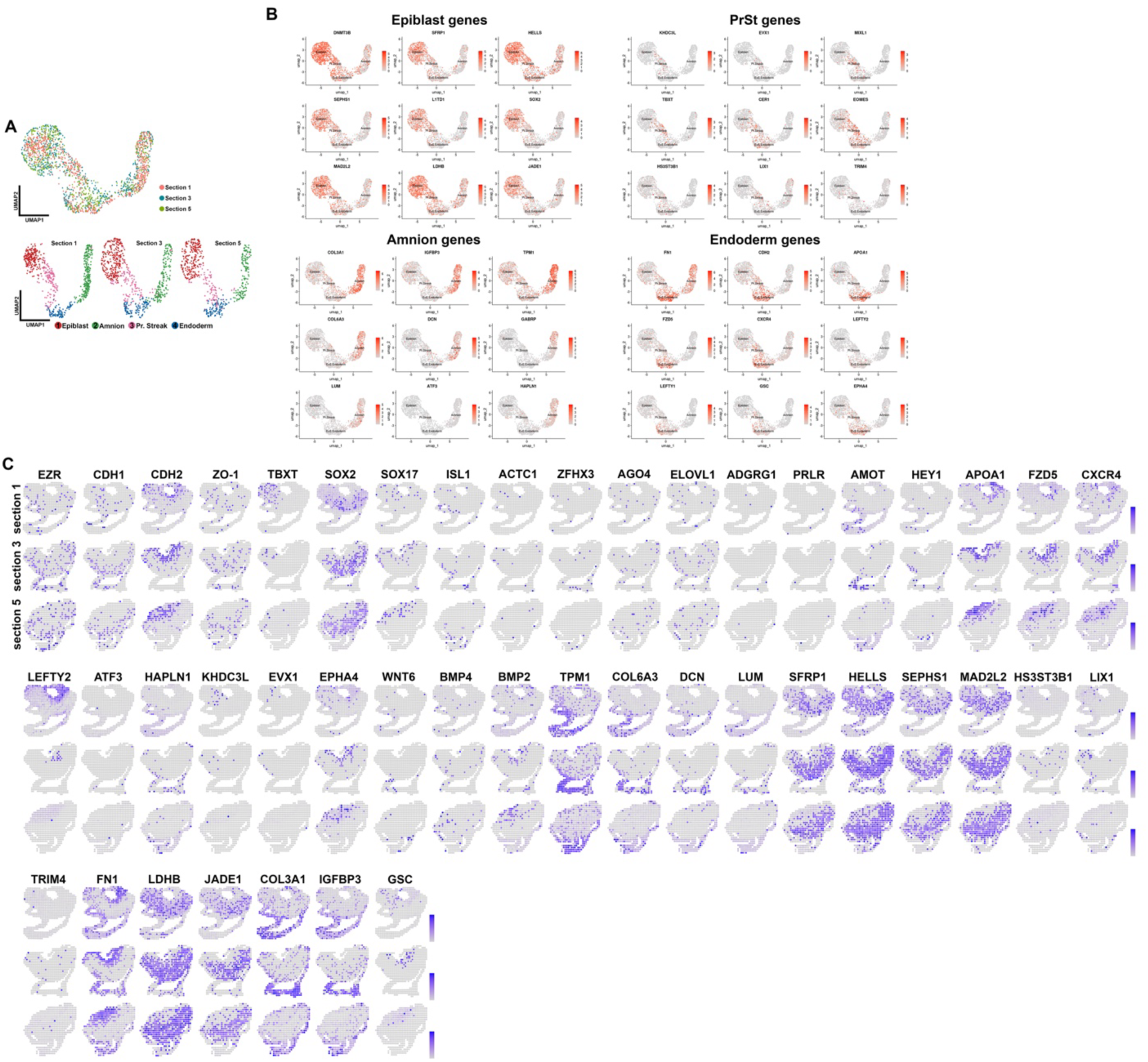
Additional information for the spatial transcriptomics. **A)** Distribution of bins (top) and bin cluster identities (bottom) for each section, projected onto the spatial transcriptomics UMAP in (Fig. 1D). **B)** Feature plots showing expression of indicated lineage markers. **C)** Expression of cluster specific DEGs superimposed onto the spatial transcriptome plot.

**Figure S4.**
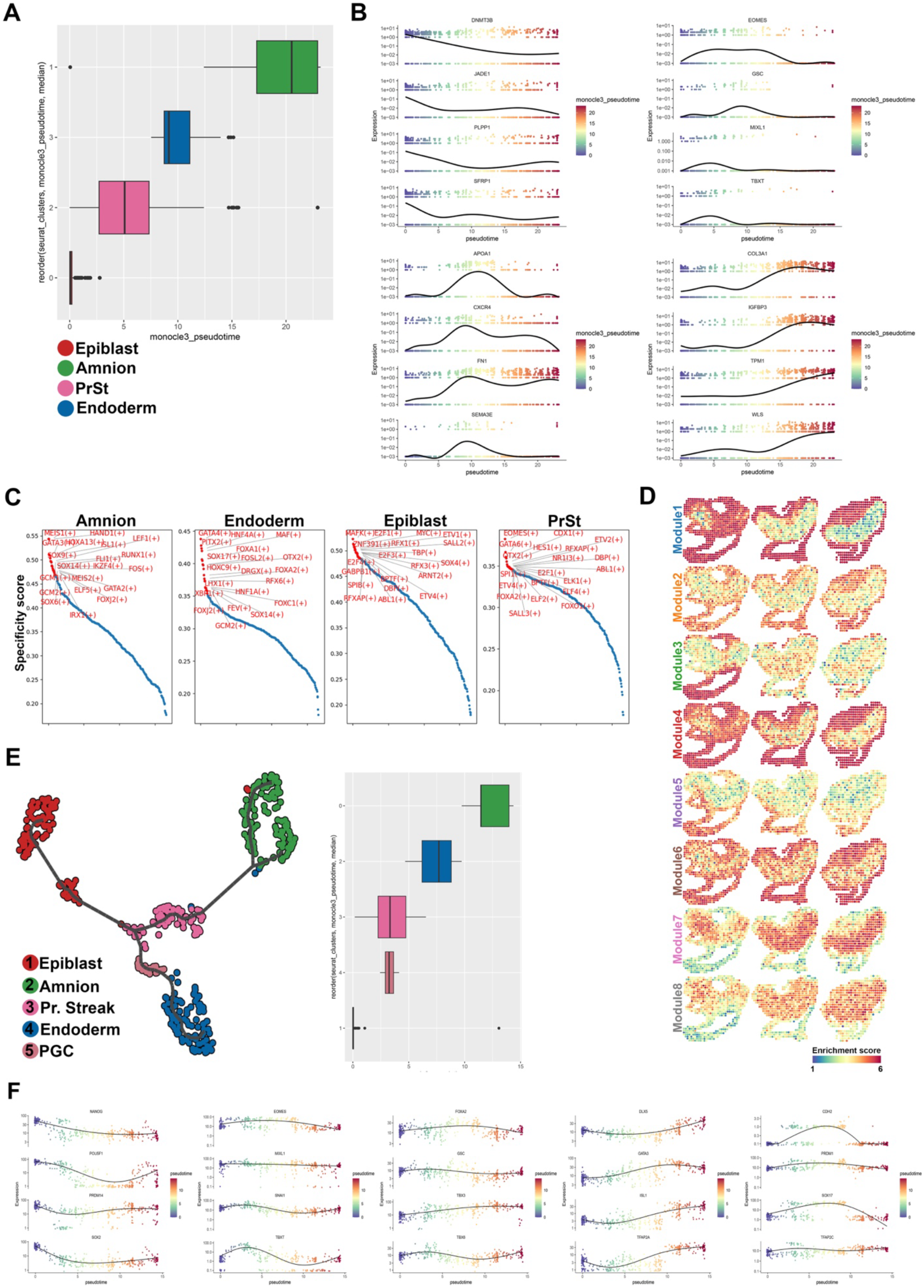
Additional information associated with pseudotime and SCENIC analyses. **A)** A boxplot showing the distribution of transcriptomic clusters along the pseudotime (in Fig. 1I). **B)** Scatterplots showing the expression dynamics of indicated genes along the pseudotime. **C)** Top 20 enriched gene regulatory networks in each transcriptomics cluster, based on SCENIC analysis (RSS plot). **D)** Enrichment scores of indicated gene regulatory network modules, based on Hotspot analysis (in Fig. 1L), superimposed onto the spatial transcriptomics plot. **E)** The spatial proteomics UMAP plot with the pseudotime trajectory (left). A box plot showing the distribution of the proteomics clusters along the pseudotime (in Fig. 1Q). **F)** Scatterplots showing the expression dynamics of indicated proteins along the proteomics pseudotime.

**Figure S5.**
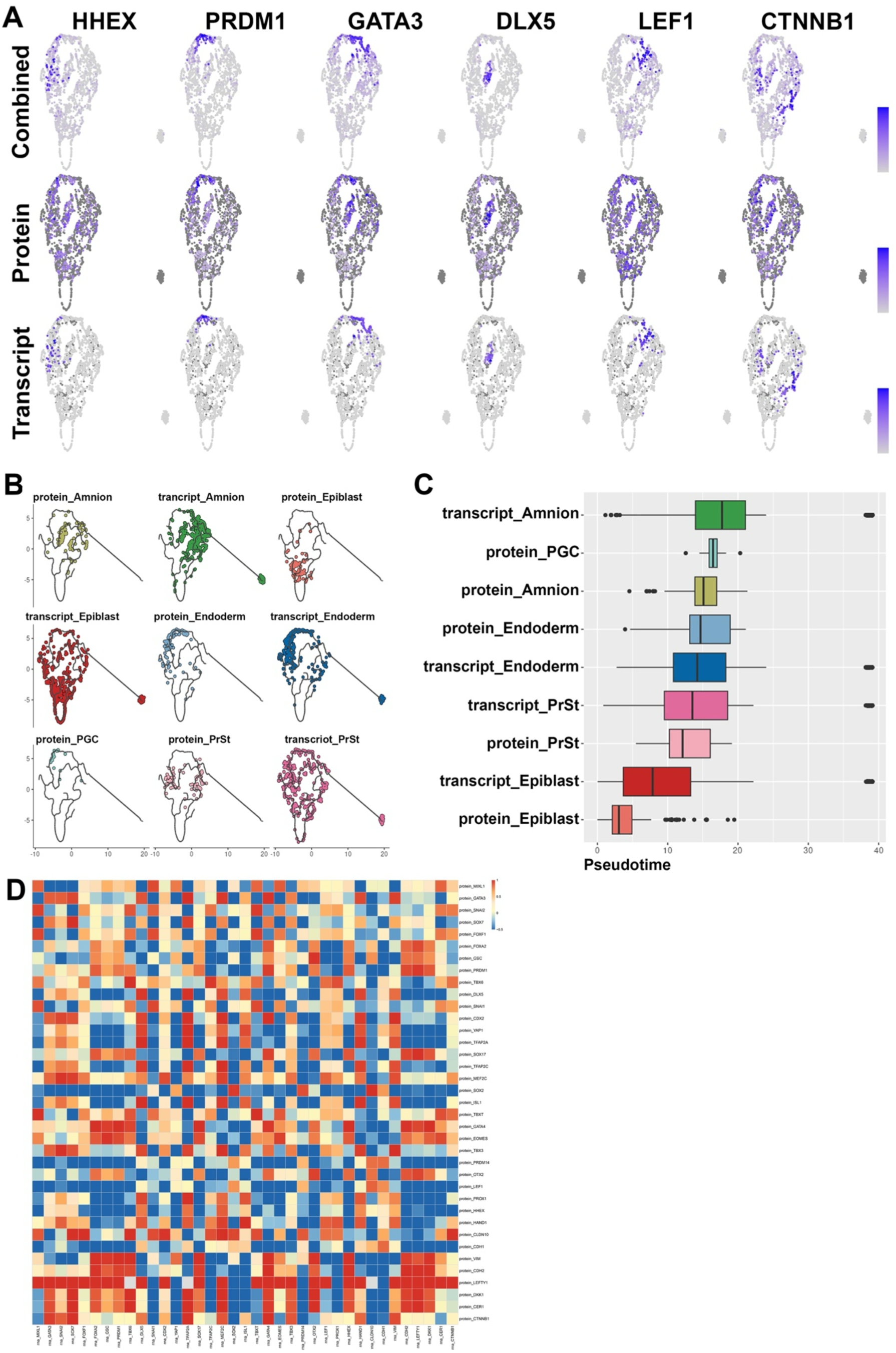
Additional information for the multiomic dataset 1. **A)** UMAP plot showing the expression of selected markers at transcript, protein, or combined (transcript and protein) levels (superimposed onto the multiomic UMAP plot in Fig. 2A). **B)** UMAP plots showing the distribution of each cell population along the multiomic pseudotime trajectory (from Fig. 2E). **C)** A box plot showing the distribution of the transcriptomic and proteomic clusters along the multiomic pseudotime. **D)** A heatmap showing Pearson correlation coefficient values for the correlation between indicated matching transcript (X-axis) and protein (Y-axis) products.

**Figure S6.**
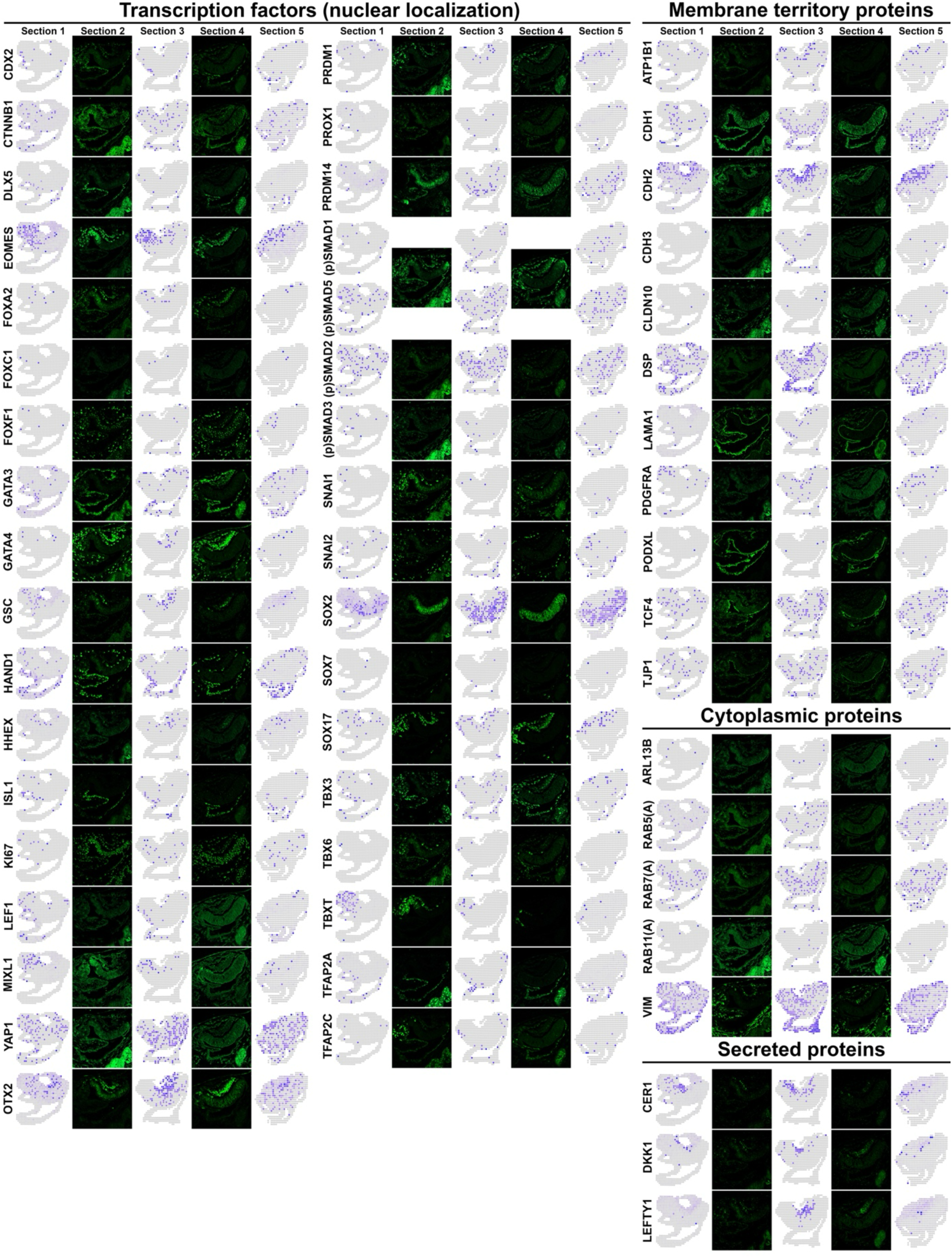
Side-by-side spatial comparisons of selected markers in the transcriptomic and proteomic datasets across five sections.

**Figure S7.**
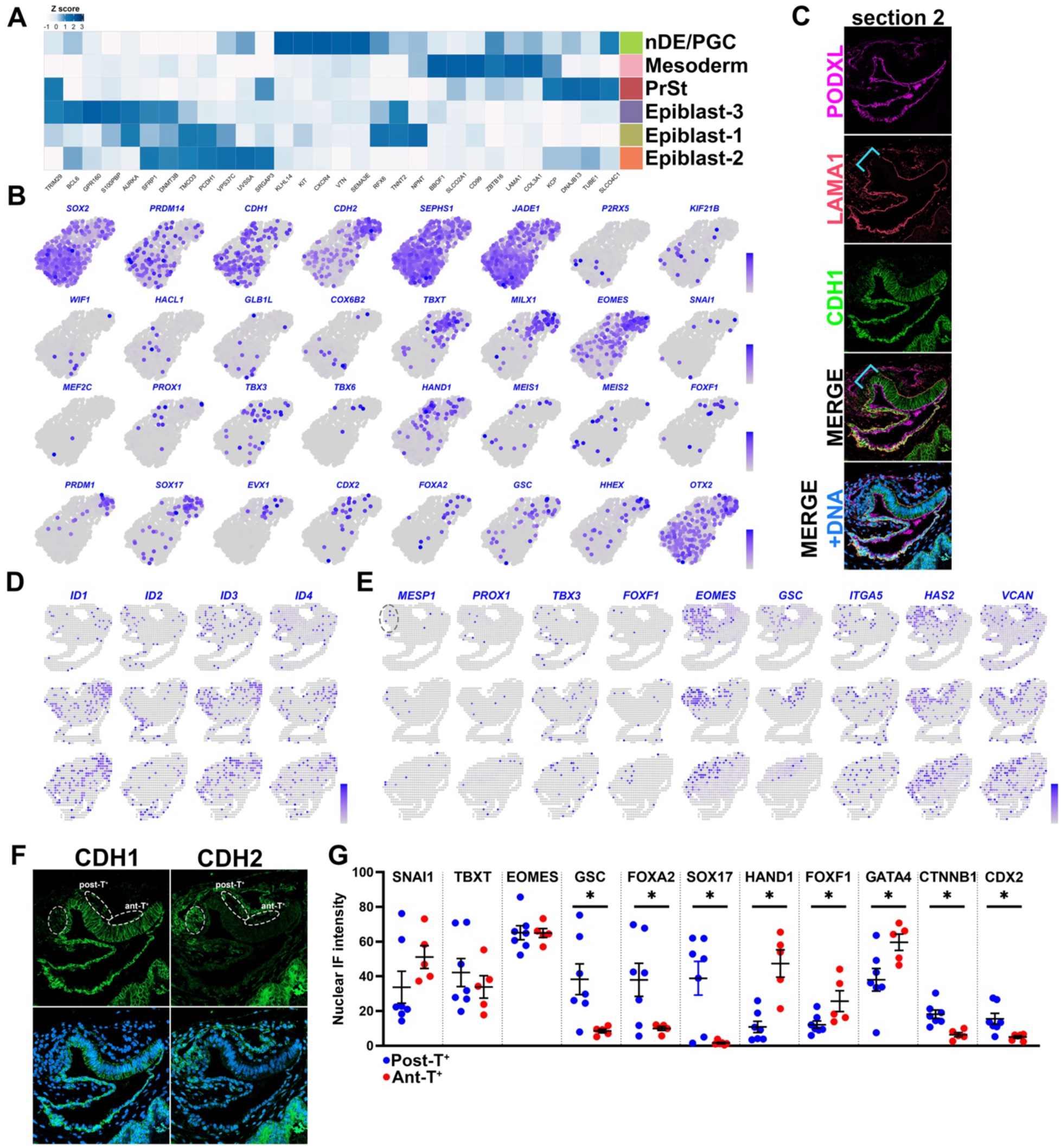
Additional information for the analysis of Epiblast and PrSt populations. **A)** A heatmap showing the normalized expression values of top 5 differentially enriched transcripts in each cell lineage. **B)** Expression of selected lineage markers superimposed onto the Epiblast-PrSt UMAP plot in Fig. 3A. **C)** Confocal IF images of the section 2, stained for indicated membrane markers. Blue bracket indicates the region in the PrSt with disorganized LAMA1. **D,E)** Expression of BMP target genes (D, *ID1-4*) and known mesoderm and allantois markers (E) superimposed onto the spatial transcriptome plot. Dotted circle in the *MESP1* plot indicates the extra-embryonic mesoderm population with PGC as well as cells that display allantois progenitor-like characteristics. **F)** Confocal IF images of the section 2, stained for CDH1 and CDH2. Light gray dashed circle indicates the extra-embryonic cell population with PGC and an early allantois-like cell population. White dashed circles indicate the posterior TBXT^+^ (post-T^+^) and anterior TBXT^+^ (ant-T^+^) cell populations. **G)** Nuclear IF intensity of indicated proteins in post-T^+^ and ant-T^+^ cell populations. Mean ± SEM shown, (Student’s t-test) * p < 0.05.

**Figure S8.**
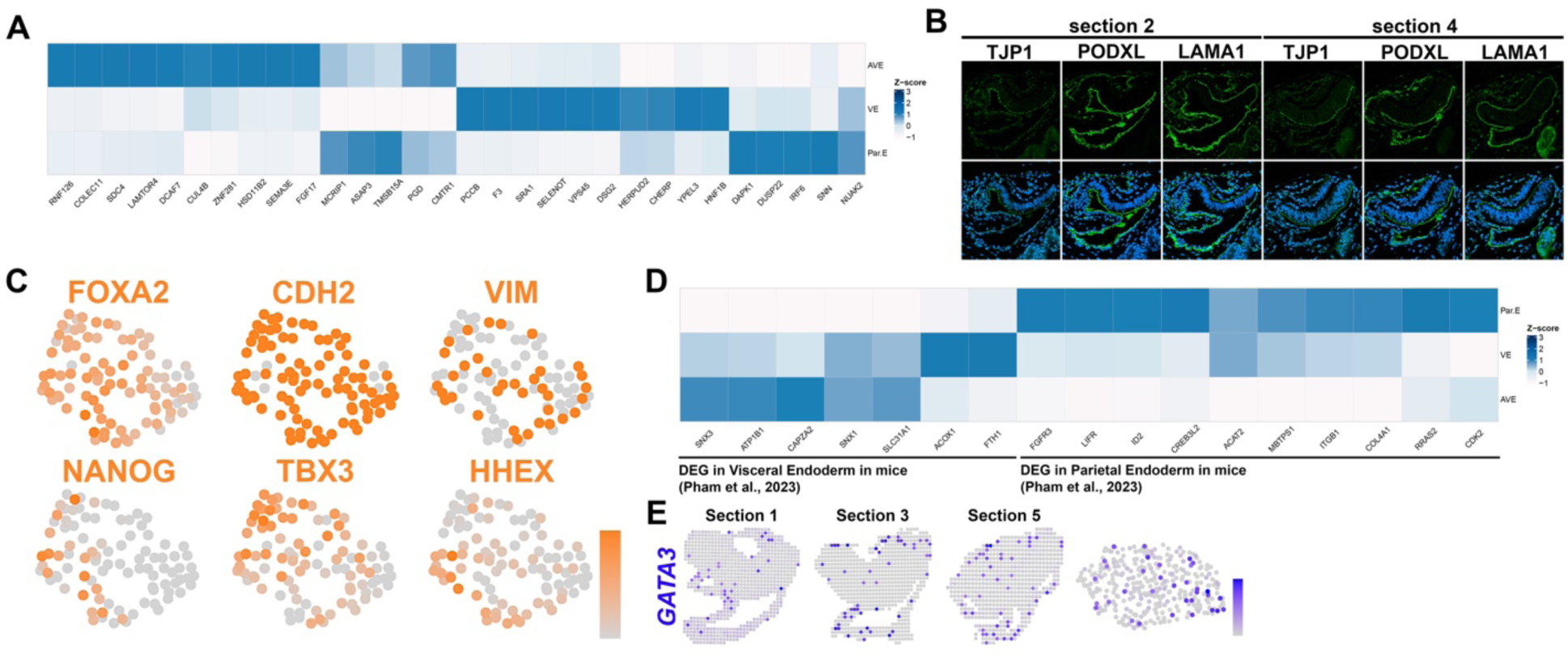
Additional information for the endoderm population. **A)** A heatmap showing the normalized expression values of the top 5 differentially enriched transcripts in each cell lineage. **B)** Confocal IF images of the CS6b embryo sections, stained for indicated markers. **C)** Expression of selected lineage specific proteins superimposed onto the endoderm spatial proteomics UMAP plot in Fig. 4C. **D)** A heatmap showing the normalized expression values of selected established VE and ParE genes in mice (markers based on Pham *et al.*, [75]). **E)** Expression of *GATA3* superimposed onto the spatial transcriptome plot and onto the endoderm UMAP plot (Fig. 4A).

**Figure S9.**
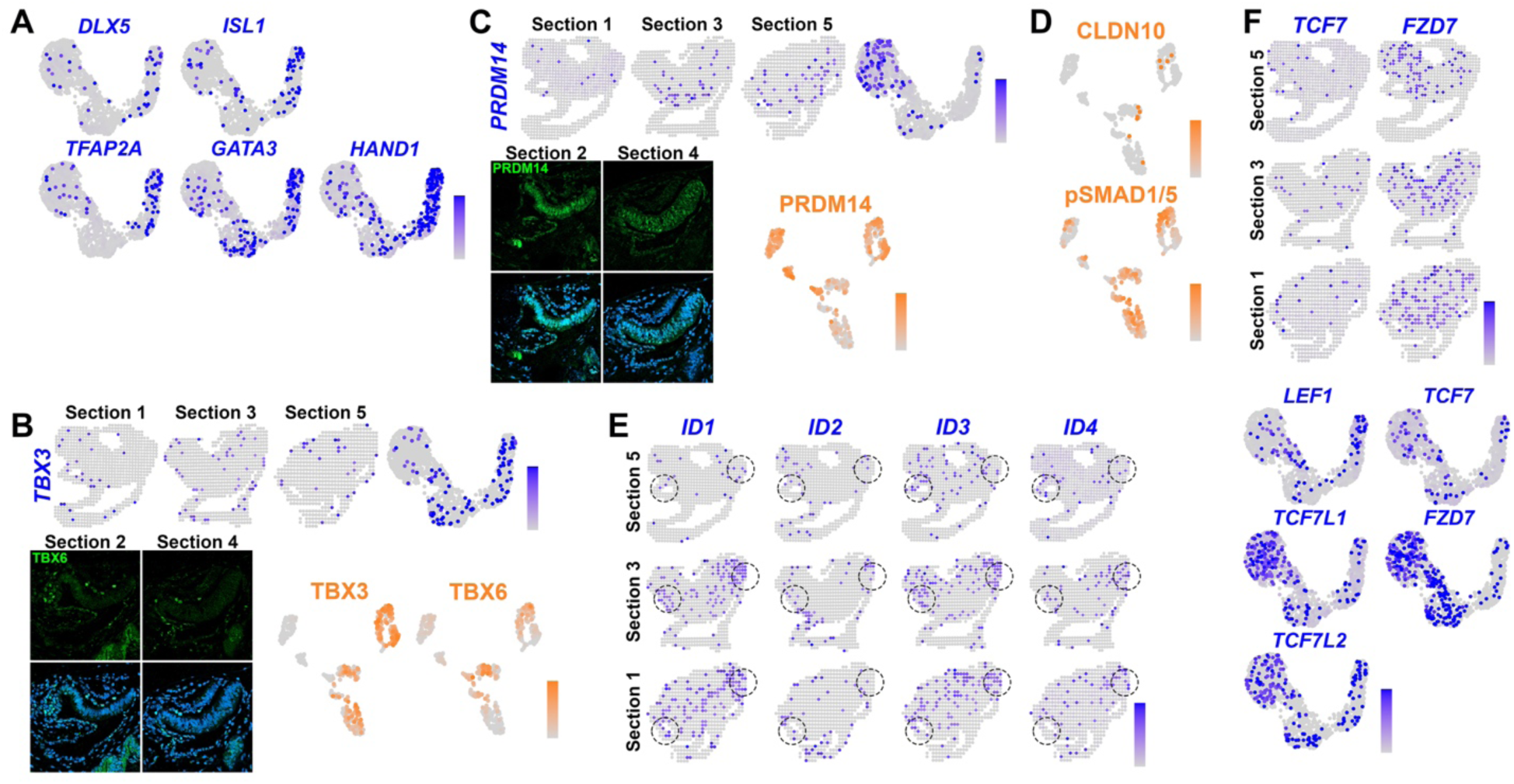
Additional information for the analysis of the amnion cell population. **A)** Expression of selected amnion markers superimposed onto the transcriptomics UMAP plot in Fig. 1D. **B)** Expression of *TBX3* superimposed onto the transcriptome plots (top, spatial and UMAP coordinates (Fig. 1D)), as well as confocal IF images of the section 2 and 4, stained for TBX6 (bottom left). Expression of TBX3 and TBX6 superimposed onto the spatial proteomic UMAP plot in Fig. 1E (bottom right). **C)** Expression analysis of PRDM14 at transcript (top) and protein (bottom) levels. **D)** Expression of CLDN10 and pSMAD1/5 superimposed onto the spatial proteomic UMAP plot (Fig. 1E). **E)** Expression of BMP target genes (*ID1-4*) superimposed onto the spatial transcriptome plot. Dotted circles indicate epiblast-amnion boundaries. **F)** Expression of WNT-dependent genes superimposed onto the spatial transcriptome plot (top) and onto the transcriptomic UMAP plot (from Fig. 1D, bottom).

**Figure S10.**
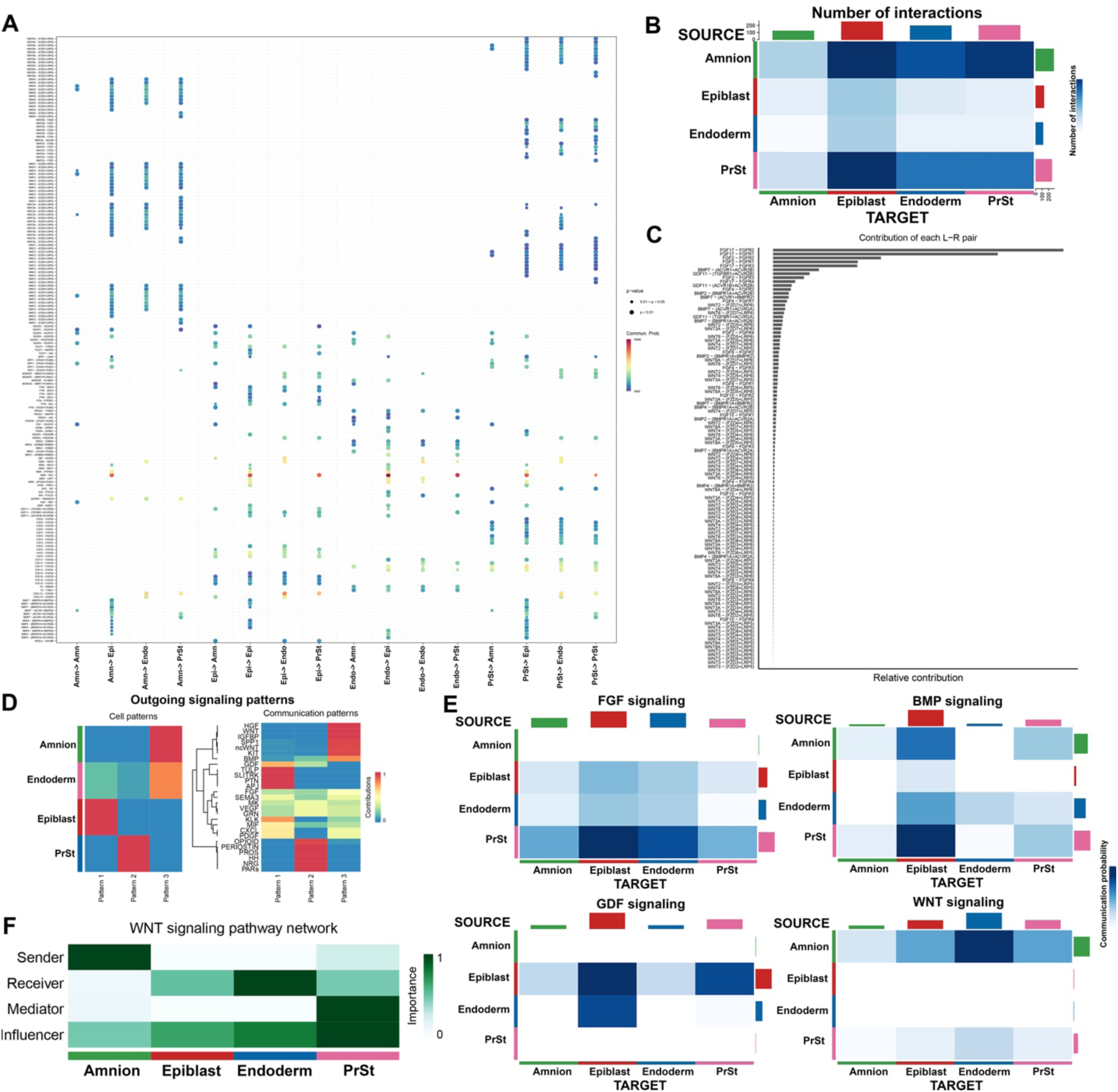
Receptor-ligand communication network analysis based on the CellChat package. **A)** A bubble plot showing the communication probability of receptor-ligand interactions (Y-axis) between different source and target cell types (X-axis). **B)** A heatmap (for signaling source and target in Y– and X-axes, respectively) showing combined significant receptor-ligand interactions across all cell types. **C)** Relative contribution of each receptor-ligand pair. **D)** Global outgoing signaling patterns for each cell type (left) or signaling pathway (right). **E)** A heatmap showing predicted signaling communication strength for indicated signaling pathways across all cell types. **F)** A heatmap showing predicted role of the WNT signaling network across all cell types.

**Figure S11.**
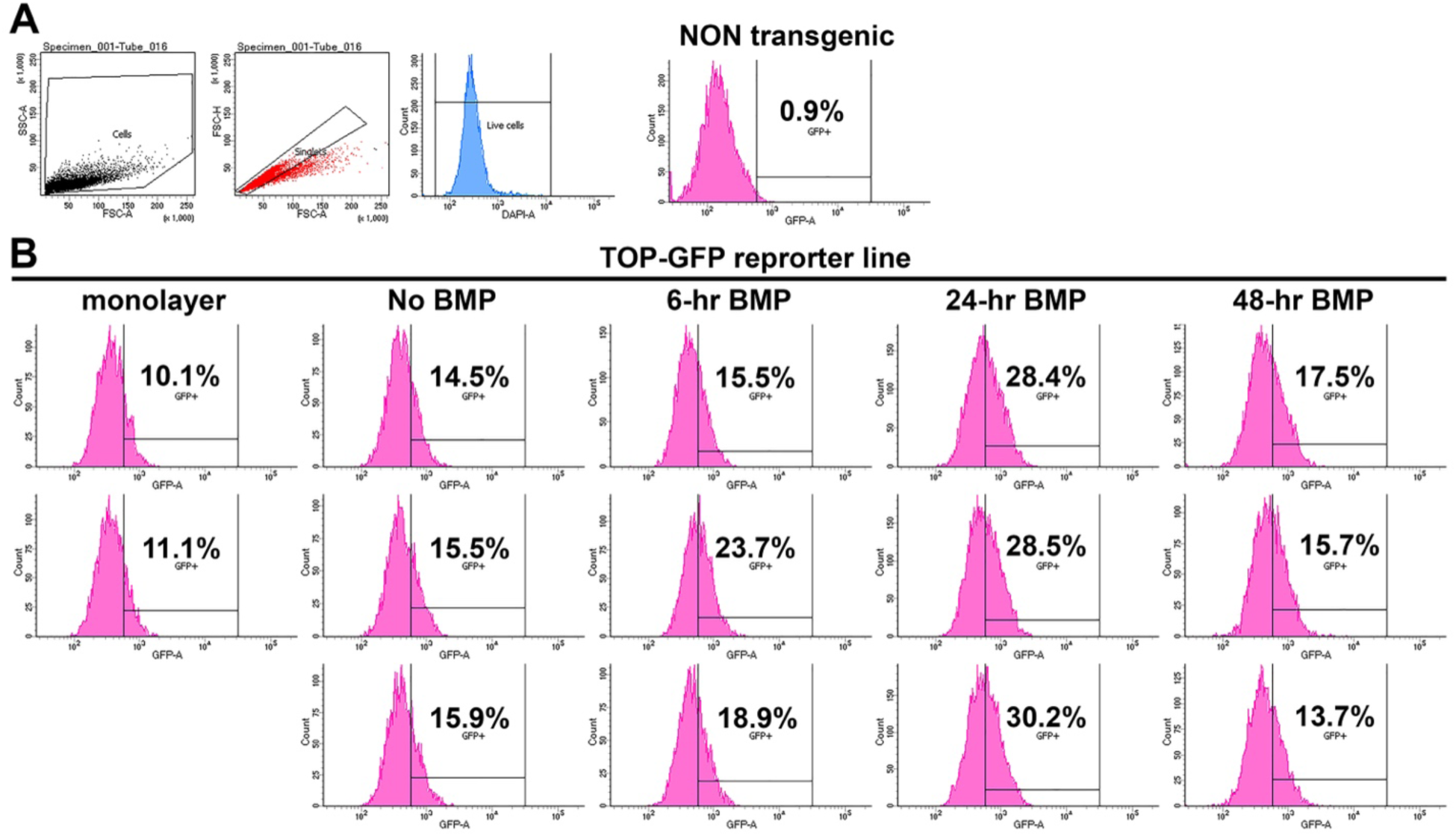
Additional flow cytometry graphs of hPSC line that expresses GFP under the control of a 6x*TCF/LEF* promoter. **A)** Gating strategy identifying singly dissociated Glass-3D^+BMP4^ cells, singlets, live cells, and GFP^+^ cell populations. **B)** Flow cytometry analysis of singly dissociated Glass-3D^+BMP4^ cells at indicated conditions, with cells gated for singlets and live cells. Percentages of GFP^+^ cells at each timepoint were calculated from the total of single and live cells.

